# A public broadly neutralizing antibody class targets a membrane-proximal anchor epitope of influenza virus hemagglutinin

**DOI:** 10.1101/2021.02.25.432905

**Authors:** Jenna J. Guthmiller, Julianna Han, Henry A. Utset, Lei Li, Linda Yu-Ling Lan, Carole Henry, Christopher T. Stamper, Olivia Stovicek, Lauren Gentles, Haley L. Dugan, Nai-Ying Zheng, Sara T. Richey, Micah E. Tepora, Dalia J. Bitar, Siriruk Changrob, Shirin Strohmeier, Min Huang, Adolfo García-Sastre, Raffael Nachbagauer, Peter Palese, Jesse D. Bloom, Florian Krammer, Lynda Coughlan, Andrew B. Ward, Patrick C. Wilson

## Abstract

Broadly neutralizing antibodies against influenza virus hemagglutinin (HA) have the potential to provide universal protection against influenza virus infections. Here, we report a distinct class of broadly neutralizing antibodies targeting an epitope toward the bottom of the HA stalk domain where HA is “anchored” to the viral membrane. Antibodies targeting this membrane-proximal anchor epitope utilized a highly restricted repertoire, which encode for two conserved motifs responsible for HA binding. Anchor targeting B cells were common in the human memory B cell repertoire across subjects, indicating pre-existing immunity against this epitope. Antibodies against the anchor epitope at both the serological and monoclonal antibody levels were potently induced in humans by a chimeric HA vaccine, a potential universal influenza virus vaccine. Altogether, this study reveals an underappreciated class of broadly neutralizing antibodies against H1-expressing viruses that can be robustly recalled by a candidate universal influenza virus vaccine.

## Introduction

Influenza viruses remain a global health problem, with antigenically drifting seasonal viruses and the constant risk of zoonotic influenza virus spillovers into humans. Antibodies against the major surface glycoprotein hemagglutinin (HA) are critical for providing protection against influenza virus infection (Ng et al., 2019). HA is divided into two domains: the globular head and the stalk. Most epitopes of the HA head are highly variable and rapidly mutate to circumvent host humoral immunity (Henry et al., 2019; Kirkpatrick et al., 2018). In contrast, the HA stalk is relatively conserved within and across influenza subtypes (Krystal et al., 1982). Antibodies against the head and the stalk both independently correlate with protection against influenza virus infection (Aydillo et al., 2020; Ng et al., 2019; Ohmit et al., 2011). Therefore, vaccine formulations that preferentially induce antibodies against conserved epitopes of the HA head and stalk domains could provide broad and potent protection against a wide array of influenza viruses.

Several broadly neutralizing epitopes have been identified on the HA of H1N1 viruses, including the receptor-binding site (RBS) and lateral patch on the HA head (Ekiert et al., 2012; Raymond et al., 2018; Whittle et al., 2011) and the broadly neutralizing (BN) epitope on the HA stalk domain (Ekiert et al., 2009; Wrammert et al., 2011). Current seasonal influenza virus vaccines poorly induce antibodies against broadly neutralizing epitopes of HA (Andrews et al., 2015; Corti et al., 2010). Therefore, new vaccine platforms that preferentially drive antibodies against these conserved epitopes are desperately needed to increase vaccine effectiveness against drifted strains and limit influenza morbidity and mortality. It is critically important to drive the humoral immune response simultaneously against multiple conserved epitopes of HA to avoid the generation of viral escape mutants. Notably, escape mutants near the lateral patch (Linderman et al., 2014; Raymond et al., 2018) and the BN stalk epitope (Park et al., 2020) have been shown to evade neutralizing antibodies at these epitopes. Hence, identification of additional broadly neutralizing epitopes of HA that can be efficiently targeted remains an important pursuit to improve vaccine effectiveness while avoiding escape mutants.

Humans are exposed to influenza viruses throughout their lifetime and reuse memory B cells (MBCs) from prior exposures to provide defense against drifted and novel strains. Seasonal influenza virus vaccines often recall MBCs targeting variable epitopes of the HA head rather than MBCs targeting conserved epitopes of HA (Andrews et al., 2015; Dugan et al., 2020). In the absence of pre-existing immunity against variable epitopes of the HA head, humans can recall MBCs targeting conserved epitopes of the HA head and stalk domains (Andrews et al., 2015). Notably, first exposure to the 2009 pandemic H1N1 virus (pH1N1) robustly recalled MBCs against conserved epitopes of the HA stalk domain (Andrews et al., 2015; Wrammert et al., 2011). Additionally, exposure to influenza viruses of zoonotic origin can recall MBCs targeting conserved epitopes of the HA stalk (Ellebedy et al., 2014; Henry Dunand et al., 2016; Nachbagauer et al., 2014).

Several leading universal influenza virus candidates function to induce antibodies specifically against the stalk domain. The chimeric HA (cHA) vaccine strategy utilizes the head domain from a zoonotic influenza virus, for which humans have little pre-existing immunity, and the stalk domain from pH1N1 (Krammer et al., 2013; Pica and Palese, 2013). A phase I clinical trial has shown cHA vaccination robustly drives protective antibodies against the stalk domain (Bernstein et al., 2020; Nachbagauer et al., 2020). In addition to the cHA vaccine strategy, several groups have generated headless HA antigens that potently induce B cells against the HA stalk in animal models while eliminating the potential of inducing B cells against the HA head domain (Impagliazzo et al., 2015; Yassine et al., 2015). The full spectrum of distinct epitopes on the HA stalk targeted by these vaccine antigens remains to be determined.

By analyzing the specificities of B cells targeting the H1 stalk through the generation of monoclonal antibodies (mAbs), we identified a class of antibodies targeting an anchor epitope of HA near the viral membrane. Antibodies targeting this epitope are broadly neutralizing across H1-expressing viruses and potently protective *in vivo*. Additionally, we showed anchor epitope targeting antibodies were recalled in humans via vaccination with both the 2009 monovalent influenza virus vaccine and by seasonal influenza virus vaccination. Furthermore, we identified that the cHA vaccine platform robustly induced antibodies against the anchor epitope. In contrast, membrane anchor targeting mAbs could not bind mini-HA, a headless HA antigen, potentially due to trimer splaying of the rHA that used a GCN4 trimerization domain. Anchor epitope targeting mAbs utilized a highly restricted repertoire and public clonotypes that encoded for two conserved motifs in the kappa chain CDR3 (K-CDR3) and heavy chain CDR2 (H-CDR2). Lastly, we identified that anchor targeting B cells are common within the human MBC pool. Together, our study reveals a novel class of broadly neutralizing antibodies against the anchor epitope, a previously unappreciated epitope. Our study additionally provides valuable insight into the binding and repertoire features of anchor epitope targeting B cells and how cHA, a potential universal influenza virus vaccine, potently induces antibodies against this epitope.

## Results

### Identification of antibodies targeting the anchor epitope

To dissect conserved HA stalk domain epitopes, we generated mAbs from acutely activated plasmablasts isolated from subjects who received licensed or experimental influenza virus vaccines or were naturally infected with pH1N1 during the 2009 pandemic (Table S1). Notably, plasmablasts found in the blood of subjects after infection or vaccination derive from pre-existing MBCs (Andrews et al., 2015), and generation of mAbs from plasmablasts allows for the dissection of how distinct influenza viruses recall pre-existing immunity. We also generated mAbs from sorted HA^+^ B cells one month following vaccination with an experimental cHA vaccine that utilized the head domain from an avian influenza virus and stalk domain from pH1N1 (Bernstein et al., 2020). We specifically focused our studies on mAbs targeting the stalk domain of H1-expressing viruses, as prior studies have shown first exposure to the 2009 pandemic H1N1 virus induce antibodies against the stalk domain (Li et al., 2012; Wrammert et al., 2011). To define antibodies as targeting the H1 stalk, mAbs were tested for binding to cH5/1, which utilizes the head domain from H5-expressing viruses and the stalk domain from the pH1N1 virus (Hai et al., 2012), and for hemagglutination inhibition (HAI) activity against pH1N1 (A/California/7/2009), a feature of head binding antibodies. MAbs that bound the cHA and that lacked HAI activity were classified as those binding the HA stalk domain. Of all mAbs tested, nearly 49% targeted the HA stalk domain, whereas 40% targeted the HA head domain (Figure S1A). To investigate what proportion were binding the BN stalk domain epitope, we competed the stalk binding mAbs from our cohorts with CR9114, a well-defined antibody targeting the BN stalk epitope (Dreyfus et al., 2012). We identified that only 21% of mAbs targeting the stalk domain had greater than 80% competition with CR9114 (Figure S1B), indicating most H1 stalk domain targeting antibodies were binding other epitopes of the HA stalk.

To investigate which epitopes the remaining 79% of mAbs were targeting on the stalk domain, we performed negative stain electron microscopy with two stalk domain binding mAbs. Both mAbs bound an epitope near the anchor of the HA stalk, towards the lower portion of the HA protomer (Figure 1A-B; Figure S1C-D). Both mAbs were oriented at an upward angle towards the epitope (Figure 1A-B), suggesting this epitope may be partially obstructed by the lipid membrane and may only be exposed for antibody binding as the HA trimers flex on the viral membrane (Benton et al., 2018). FISW84, a recently identified anchor binding mAb, (Benton et al., 2018), overlap with both 047-09 4F04 and 241 IgA 2F04 (Figure 1C), suggesting this epitope is a common target of stalk binding antibodies. The footprint of several BN stalk epitope binding mAbs (CR9114 and FI6v3) did not overlap with those of 047-09 4F04 and 241 IgA 2F04 (Figure 1D; Figure S1E), indicating the anchor epitope and the BN stalk epitope are distinct epitopes on the HA stalk. To understand what proportion of stalk binding mAbs were binding to the anchor epitope, we competed 047-09 4F04 mAb with the remaining stalk binding mAbs that did not compete with CR9114. In total, we identified 50 distinct mAbs that competed for binding to the anchor epitope from a total of 21 subjects (Table S2) and accounted for 28% of all stalk mAbs identified (Figure 1E-F). Together, these data indicate that the anchor epitope is a common target of antibodies binding the H1 stalk domain.

**Figure 1:**
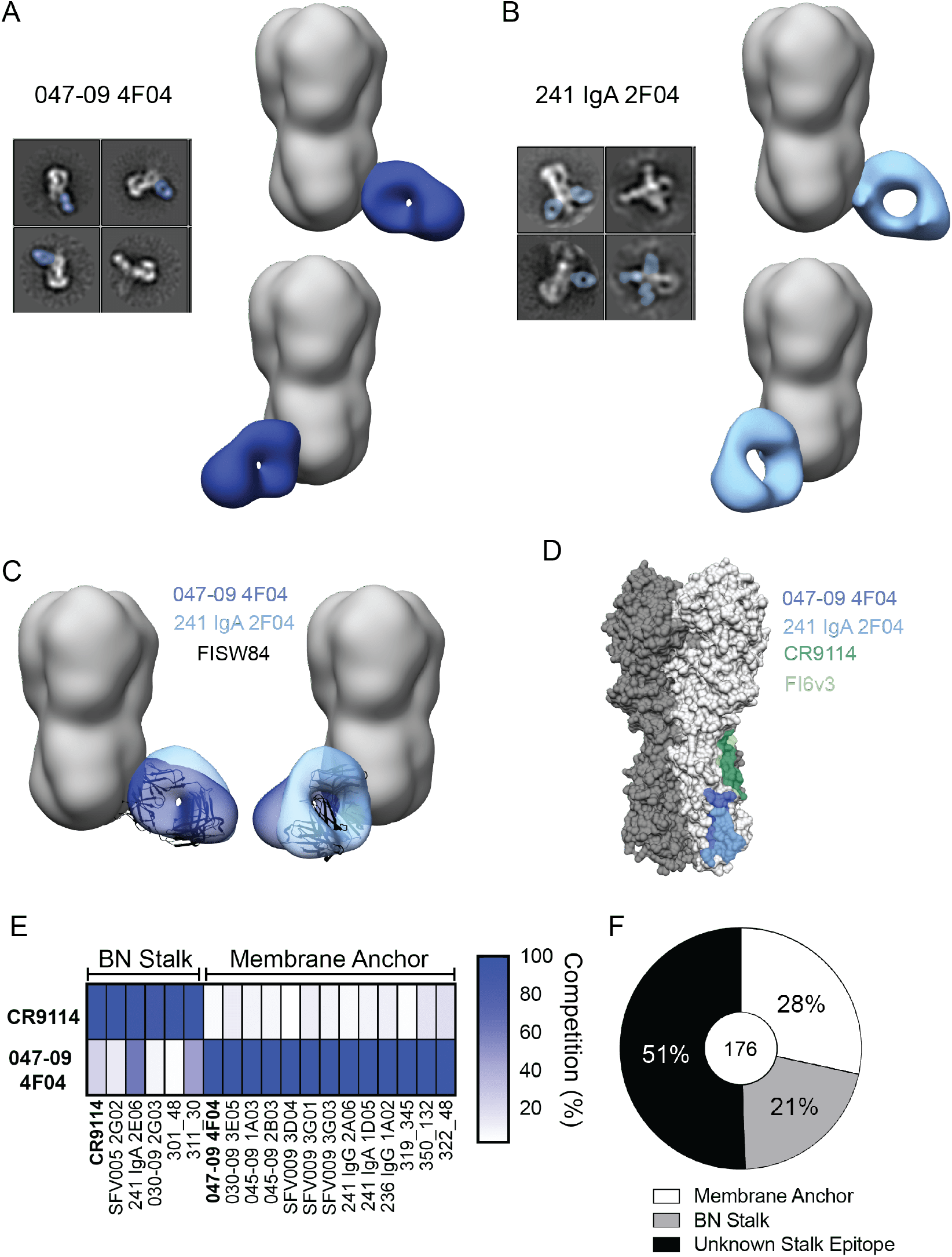
The anchor epitope is a common target of HA stalk binding antibodies. **(A**-**B)**, Negative stain EM 2D class averages and 3D reconstructions of negative stain EM of 047-09 4F04 Fab (**A**) and 241 IgA 2F04 Fab (**B**) binding to A/California/4/2009 HA. (**C**) Overlay of 047-09 4F04, 241 IgA 2F04, and FISW84 (PDB: 6HJQ) Fabs binding the anchor epitope of A/California/04/2009 HA. (**D**) Binding footprints of 047-09 4F04, 241 IgA 2F04, CR9114, and FI6v3 on A/California/04/2009 HA. (**E**) Competition of stalk binding mAbs with CR9114 or 047-09 4F04. (**F**) Proportion of mAbs binding to the anchor epitope, the BN stalk epitope, or an unknown stalk epitope based on competition with 047-09 4F04 or CR9114. Number in the center of the pie graph represents the number of mAbs tested. See also **Figure S1 and Table S2**.

### Antibodies binding the anchor are broadly reactive amongst H1 viruses

As the stalk domain is conserved amongst influenza viruses, we next determined the viral binding breadth of antibodies targeting the anchor epitope. Anchor mAbs were broadly reactive amongst H1-expressing viruses, including a swine origin H1N2 virus, but rarely cross-reacted with other influenza subtypes (Figure 2A; Figure S2A-B), as often occurs for antibodies targeting the BN stalk epitope (Figure 2A; Figure S2B). While highly conserved amongst H1 viruses, the anchor epitope was poorly conserved across divergent group 1 viral subtypes (Figure S2C). Anchor epitope targeting mAbs had nearly a 2-fold higher affinity for pH1N1 virus than mAbs targeting the BN stalk epitope (Figure 2B). Because the anchor epitope is partially obstructed by the lipid membrane, we next tested whether anchor binding mAbs had reduced affinity for whole virus relative to recombinant HA (rHA). MAbs binding the anchor epitope and the BN stalk epitope both exhibited reduced affinity for the whole virus (A/California/7/2009) relative to rHA from the same virus, whereas mAbs targeting the HA head domain had similar affinity for whole virus and rHA (Figure S2D). These data indicate that antibodies targeting the anchor epitope are broadly reactive amongst H1-expressing viruses.

**Figure 2:**
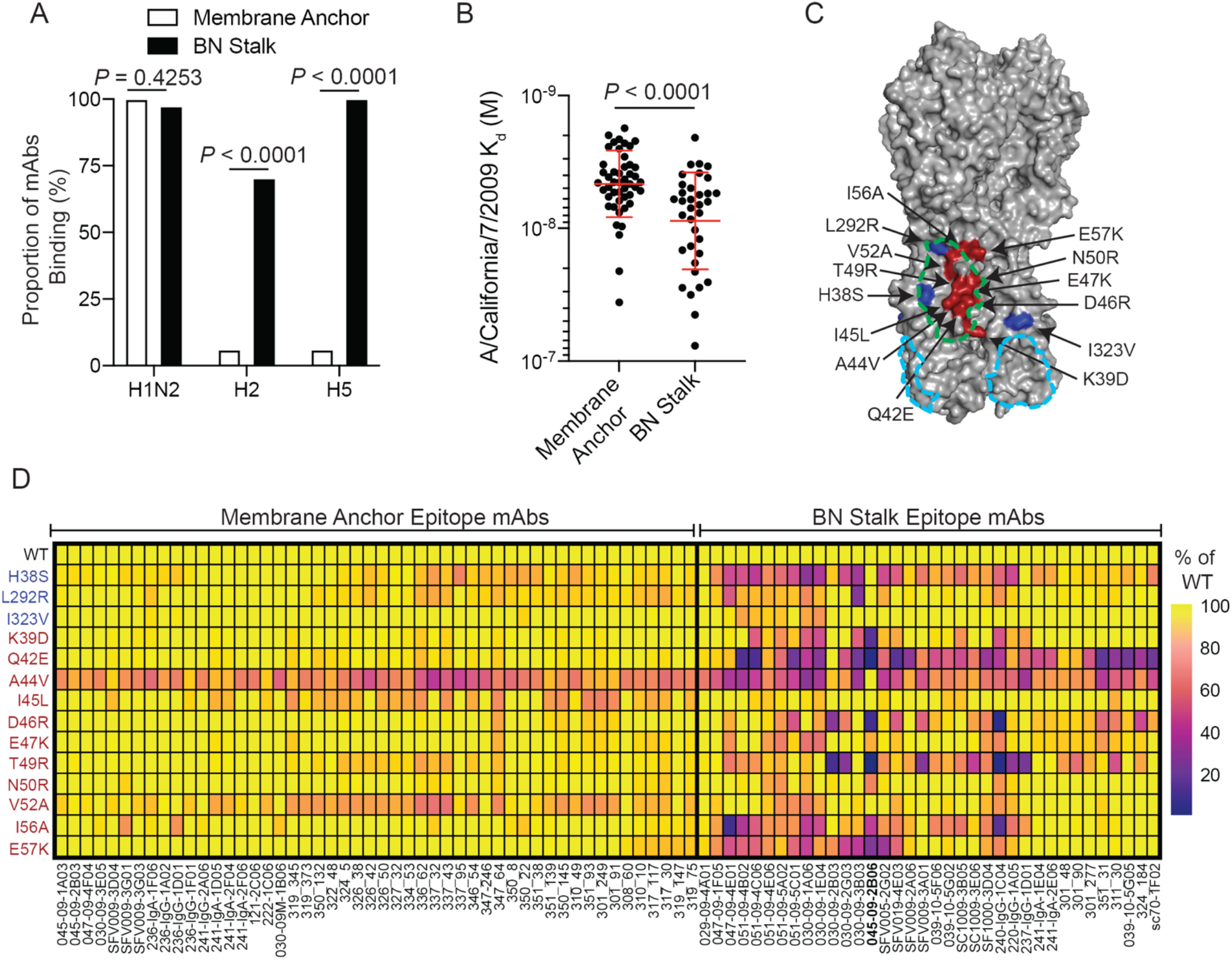
Anchor epitope targeting antibodies are broadly reactive amongst H1 expressing viruses. (**A**) Proportion of anchor epitope and BN stalk epitope targeting mAbs binding to other group 1 subtypes. (**B**) Apparent affinity of anchor and BN stalk binding mAbs to A/California/7/2009 virus. Data are represented as mean ± S.D. (**C**-**D**) Anchor and BN stalk binding mAbs were tested for binding to A/California/7/2009 HA with naturally occurring mutations and experimentally determined mutations induced by 045-09 2B06, a BN stalk epitope binding mAb. (**C**) Location of mutations modeled on A/California/04/2009 HA (PDB: 4JTV). Residues in blue are located on HA1 and residues in red are located on HA2. Outlines represent binding footprints of 047-09 4F04 (sky blue) and CR9114 (green). (**D**) Heatmap of mAb binding to WT and mutant HAs shown as the proportion of signal relative to mAb binding to the WT HA. Data in **A** were analyzed by Fisher’s Exact tests and **B** were analyzed by unpaired non-parametric Mann-Whitney test. See also **Figure S2**, **Figure S3, Table S3**.

### Antibodies targeting the anchor epitope maintain binding to HA mutants in the stalk domain

H1N1 viruses have acquired several mutations within the HA stalk domain, likely due to antibody selective pressures or to increase stability (Cotter et al., 2014). To understand whether these mutations have affected antibody binding to the anchor epitope, we screened mAbs against naturally occurring mutants and experimentally identified viral escape mutants of BN stalk epitope binding mAbs (Figure 2C-D; Figure S3A-B; Table S3). Anchor epitope binding mAbs were mostly unaffected by the mutants tested, whereas most of the mAbs targeting the BN stalk epitope were affected by at least one mutant, notably Q42E mutation in HA2 (Figure 2D). Regardless of mAb specificity, most antibodies had reduced binding to A44V of HA2, which was recently shown to preferentially grow in the presence of mAbs against the BN stalk epitope (Park et al., 2020). While A44 is distant from the anchor epitope, the A44V mutation was shown to affect the conformation of the HA stalk (Park et al., 2020) and could explain the broad reduction of HA binding by antibodies targeting either the anchor epitope or the BN stalk epitope. Furthermore, A/Michigan/45/2015 acquired mutations at S124N and E172K of HA2 (Clark et al., 2017), which lay near the binding footprint of 047-09 4F04 and 241 IgA 2F04 (Figure S3C). Despite this, mAbs targeting the anchor epitope or BN stalk epitope bind A/California/7/2009 (S124, E172) with nearly identical affinity as they do to A/Michigan/45/2015 (N124, K172; Figure S3D-E), indicating these mutations were likely not driven by selective pressures of mAbs targeting these residues. Together, these data indicate that known mutations within the HA stalk largely do not affect the binding of antibodies to the anchor epitope.

### Antibodies targeting the anchor epitope are broadly neutralizing against H1N1 viruses and are potently protective *in vivo*

We next determined whether mAbs targeting the anchor epitope were neutralizing. All mAbs targeting the anchor epitope and the BN stalk epitope were neutralizing against the pH1N1 virus (A/California/7/2009) and had similar neutralizing potency relative to mAbs binding the BN stalk epitope (Figure 3A). Furthermore, anchor targeting mAbs were broadly neutralizing against historical and recent H1N1 viruses, as well as a swine H1N2 (A/swine/Mexico/AVX8/2011) virus (Figure 3B). Together, these data indicate that antibodies against the anchor epitope and other broadly neutralizing epitopes could work in tandem to be potently neutralizing against antigenically drifted and shifted H1-expressing viruses.

**Figure 3:**
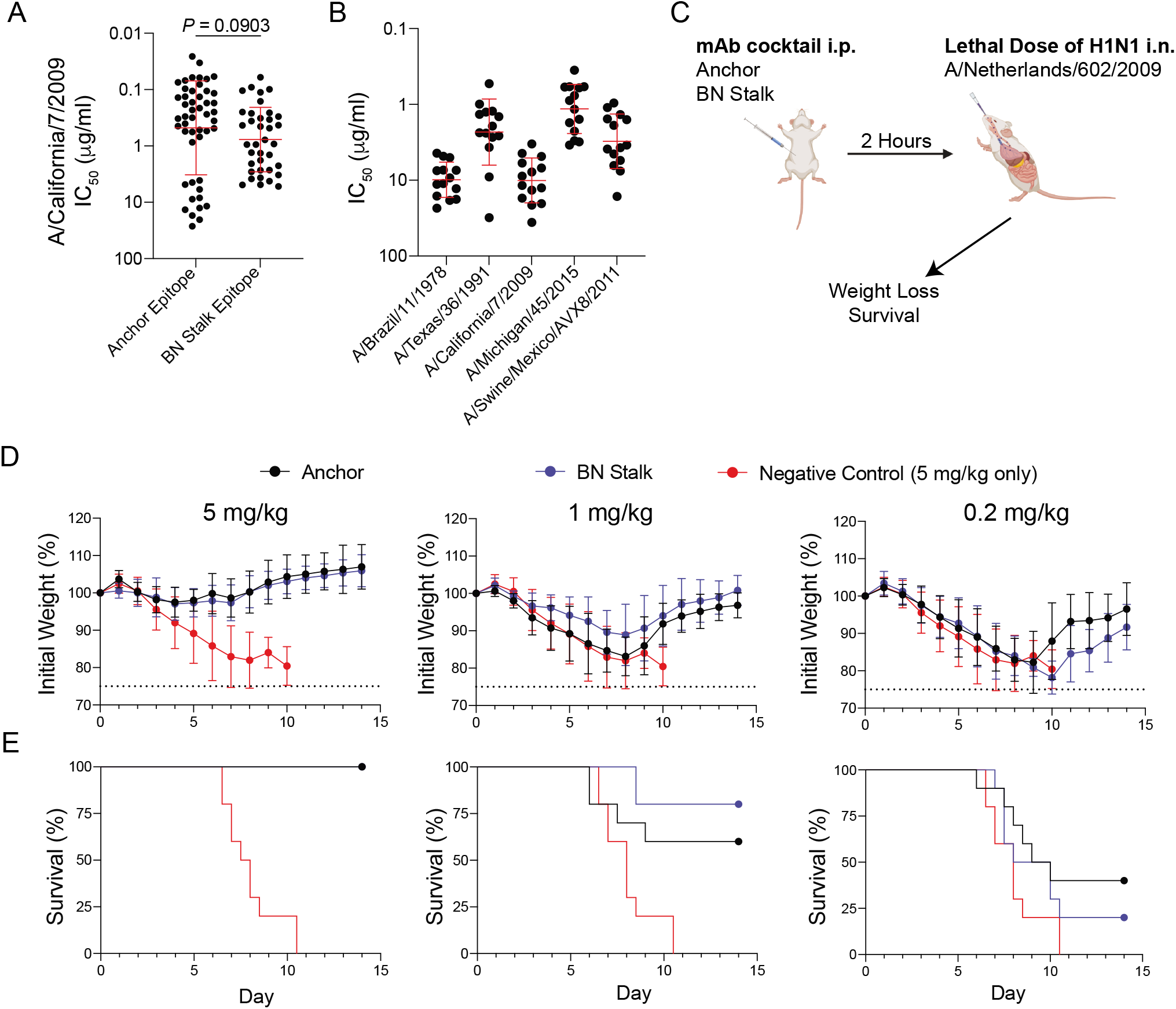
Anchor epitope targeting mAbs are broadly neutralizing amongst H1 viruses and potently protective *in vivo*. (**A**) Neutralization potency of mAbs binding the anchor or BN stalk epitope against A/California/7/2009 H1N1. (**B**) Neutralization potency of anchor epitope binding mAbs against H1-expressing viruses. (**C**-**E**) Mice were prophylactically administered i.p. a cocktail of mAbs (n=5 mAbs/cocktail) against the anchor epitope or BN stalk epitope, or an anthrax specific antibody. Mice were infected 2 hours later with 10 LD50 of A/Netherlands/602/2009 H1N1. (**C**) Experiment design. Weight loss (**D**) and survival (**E**) of mice in each treatment group. N=10 mice per treatment group and are pooled from two independent experiments. Data in **A**, **B**, and **D** are represented as mean ± S.D. Data in **A** were analyzed by unpaired non-parametric Mann-Whitney test. See also **Table S4**.

To test whether mAbs targeting the anchor epitope were protective *in vivo*, we prophylactically administered a cocktail of 5 mAbs targeting the anchor epitope or the BN stalk epitope to mimic a polyclonal response against these epitopes and infected mice with a lethal dose of a mouse-adapted pH1N1 virus (A/Netherlands/602/2009; Figure 3C). Mice that received either cocktail lost a similar amount of weight and experienced similar mortality (Figure 3D-E). Notably, mice were completely protected at 5 mg/kg of mAb cocktail, whereas only 60-80% of animals survived at 1 mg/kg (Figure 3E). Together, these data reveal mAbs targeting the anchor epitope are protective *in vivo* and could provide broad protection against H1-expressing viruses.

### Anchor epitope targeting antibodies are induced by seasonal influenza virus vaccines

Antibodies induced by influenza vaccination are biased toward variable epitopes of the HA head (Figure 4A). However, novel exposure to the 2009 pH1N1 virus robustly recalled MBCs targeting the conserved epitopes of the HA head and stalk domains, likely because subjects had low pre-existing antibody titers against the variable epitopes of the HA head (Andrews et al., 2015; Guthmiller et al., 2020; Li et al., 2012; Wrammert et al., 2011). In contrast, subjects that have been repeatedly exposed to the pH1N1 virus tend to recall MBCs targeting the variable epitopes of the HA head (Guthmiller et al., submitted for publication). Consistent with this, subjects that received the 2009 monovalent influenza virus (MIV) vaccine robustly induced a plasmablast response against the stalk domain (38%), as determined by generated mAbs, whereas only 15% of mAbs isolated from subjects that received the seasonal vaccine 2014 quadrivalent influenza virus vaccine (QIV) targeted the stalk domain (Figure 4A). Most subjects in each cohort had at least one stalk domain-targeting mAb isolated, although the frequency of stalk domain-binding mAbs per subject was higher in the subjects that received the 2009 MIV relative to subjects in the 2014 QIV cohort (Figure S4A-B). When broken down by the specific stalk epitopes targeted, nearly 40% of stalk binding mAbs isolated targeted the BN stalk epitope (Figure 4B). A larger proportion of mAbs (57%) targeting the anchor epitope were isolated from subjects that received the 2014 QIV relative to those subjects that received the 2009 MIV (16%; Figure 4B). Anchor epitope binding mAbs were detected in two out of six subjects in the 2014 QIV cohort (33.3%) and four out of eleven subjects in the 2009 MIV cohort (36%; Figure 4C), demonstrating that this epitope is commonly targeted after influenza virus vaccinations.

**Figure 4:**
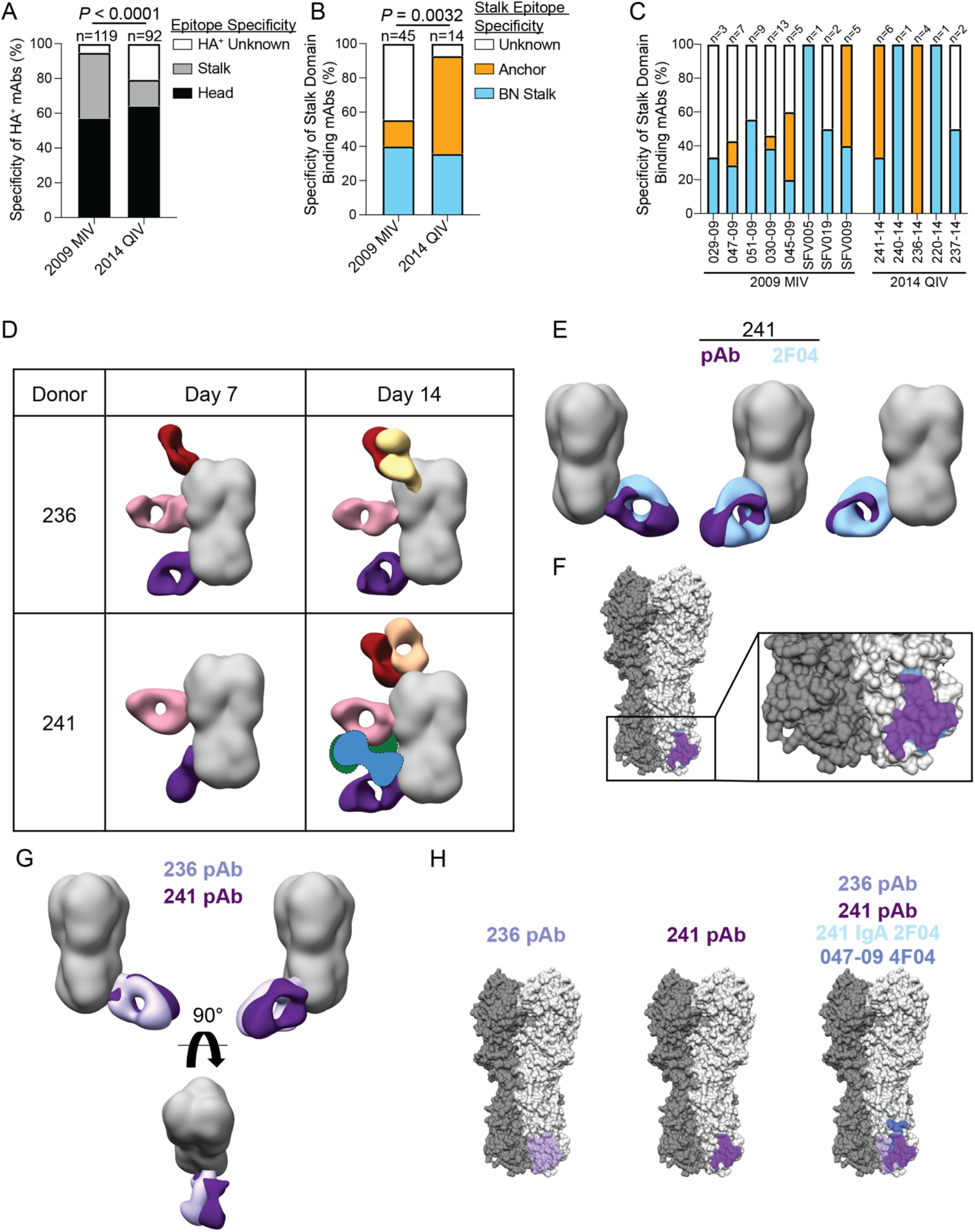
Anchor epitope targeting B cells are induced by licensed influenza virus vaccines. (**A-C**) MAbs were generated from plasmablasts isolated 7 days after influenza virus vaccination with the 2009 MIV and the 2014 QIV. (**A**) Domain binding of HA^+^ mAbs. (**B**-**C**) Epitope specificity of stalk domain binding mAbs by vaccine cohort (**B**) and by subjects (**C**). (**D**-**F**) EMPEM of serum collected at day 7 and 14 following 2014 QIV in subjects 236 and 241 binding to A/Michigan/45/2015 HA. (**D**) Summary of pAbs at day 7 and d14. (**E**) Overlap of 241 IgA 2F04 fab and pAb binding anchor epitope from subject 241. (**F**) Binding footprint of 241 IgA 2F04 (sky blue) and pAb from subject 241 (purple). (**G**) Overlap of anchor epitope binding pAbs from subjects 236 (lavender) and 241 (purple). (**H**) Binding footprint of pAbs from subjects 236 (lavender) and 241 (purple) relative to 241 IgA 2F04 and 047-09 4F04. Data in **A** and **B** were analyzed using Chi-square tests. See also **Figure S4**.

To confirm that anchor epitope targeting mAbs generated from plasmablasts were representative of the serum antibody response, we performed electron microscopy polyclonal epitope mapping (EMPEM) to dissect the targets of the polyclonal serum antibody response mounted by subjects 236 and 241 from the 2014 QIV cohort. Both subjects had detectable antibodies targeting the anchor epitope at days 7 and 14 post vaccination, whereas only subject 241 had detectable antibodies against the BN stalk epitope at day 14 (Figure 4D-E; Figure S4C-D). Notably, subject 241 had more complexes with antibodies targeting the anchor than the BN stalk epitope (Figure S4D), suggesting this subject more readily mounted an antibody response against the anchor epitope. Comparison of anchor epitope binding polyclonal antibodies (pAbs) identified in subjects 236 and 241 revealed the 241 IgA 2F04 mAb strongly overlapped with the 241 pAb from the same donor (Figure 4E-F) while the 236 pAb sat slightly anterior to the HA trimer, similar to 047-09 4F04 (Figure 4G-H; Figure S4E). Together, these data indicate influenza virus vaccination can recall MBCs targeting the anchor epitope.

### The cHA universal influenza virus vaccine candidate robustly induced antibodies against the anchor epitope

Several experimental universal influenza virus vaccine candidates currently being tested in clinical trials are intended to induce antibodies against the stalk domain. Notably, a cHA vaccine platform was shown to specifically induce antibodies against the stalk domain (Bernstein et al., 2020; Nachbagauer et al., 2020). To investigate whether subjects who received the cHA vaccine mounted an antibody response against the anchor epitope and the BN stalk epitope, we adapted the competition ELISA to detect serum antibody responses that could compete for binding with 047-09 4F04 and CR9114, respectively. Subjects enrolled in the cHA clinical trial received a prime-boost regimen of cHA, with the prime being a cH8/1 inactivated influenza virus vaccine with an adjuvant (IIV+AS03) or cH8/1 live-attenuated influenza virus (LAIV) followed by a boost 3 months later with a cH5/1 IIV with or without adjuvant (Figure 5A). On the prime vaccination, subjects that received the cH8/1 IIV+AS03 had a dramatic increase in serum antibody responses against the anchor epitope and BN stalk epitope relative to the placebo group (Figure 5B) and had a 3-fold increase in antibodies binding the anchor epitope over the day 0 time point (Figure S5A-B). However, these titers drastically dropped after 3 months post-prime (Figure 5B). Subjects that received the cH8/1 LAIV did not have an increase in serum antibody responses against either the anchor epitope or the BN stalk epitope (Figure 5B; Figure S5A-B). After the cH5/1 boost, subjects in the LAIV/IIV+AS03 group dramatically increased antibody titers against both the anchor epitope and BN stalk epitope, whereas subjects in the LAIV/IIV cohort with no adjuvant did not have a substantial increase in serum antibodies compared to the placebo cohort (Figure 5B). Subjects that received the cH8/1 IIV+AS03 followed by the cH5/1 IIV+AS03 also boosted antibody responses against both the anchor and BN stalk epitopes compared to the placebo controls (Figure 5B), although the fold-change in titers was not statistically greater than the placebo (Figure S5A-B). Furthermore, only subjects in the LAIV/IIV+AS03 cohort had a significant fold-increase in antibodies targeting both the anchor and BN stalk epitopes relative to pre-boost titers (Figure S5A-B). At a 1-year time point, subjects within the LAIV/IIV+AS03 and the IIV+AS03/IIV+AS03 had a significant decrease in serum antibodies against the anchor epitope and subjects within the LAIV/IIV+AS03 cohort had a significant decrease in serum antibodies against the BN stalk epitopes (Figure S5C-D). Furthermore, we identified and generated anchor epitope targeting mAbs from acutely activated plasmablasts isolated from one subject that received the cH8/1 IIV+AS03 prime and one subject that received the cH5/1 IIV+AS03 boost (Table S1 and Table S2). Together, these data indicate that the cHA vaccine strategy can robustly induce antibodies against the anchor epitope.

**Figure 5:**
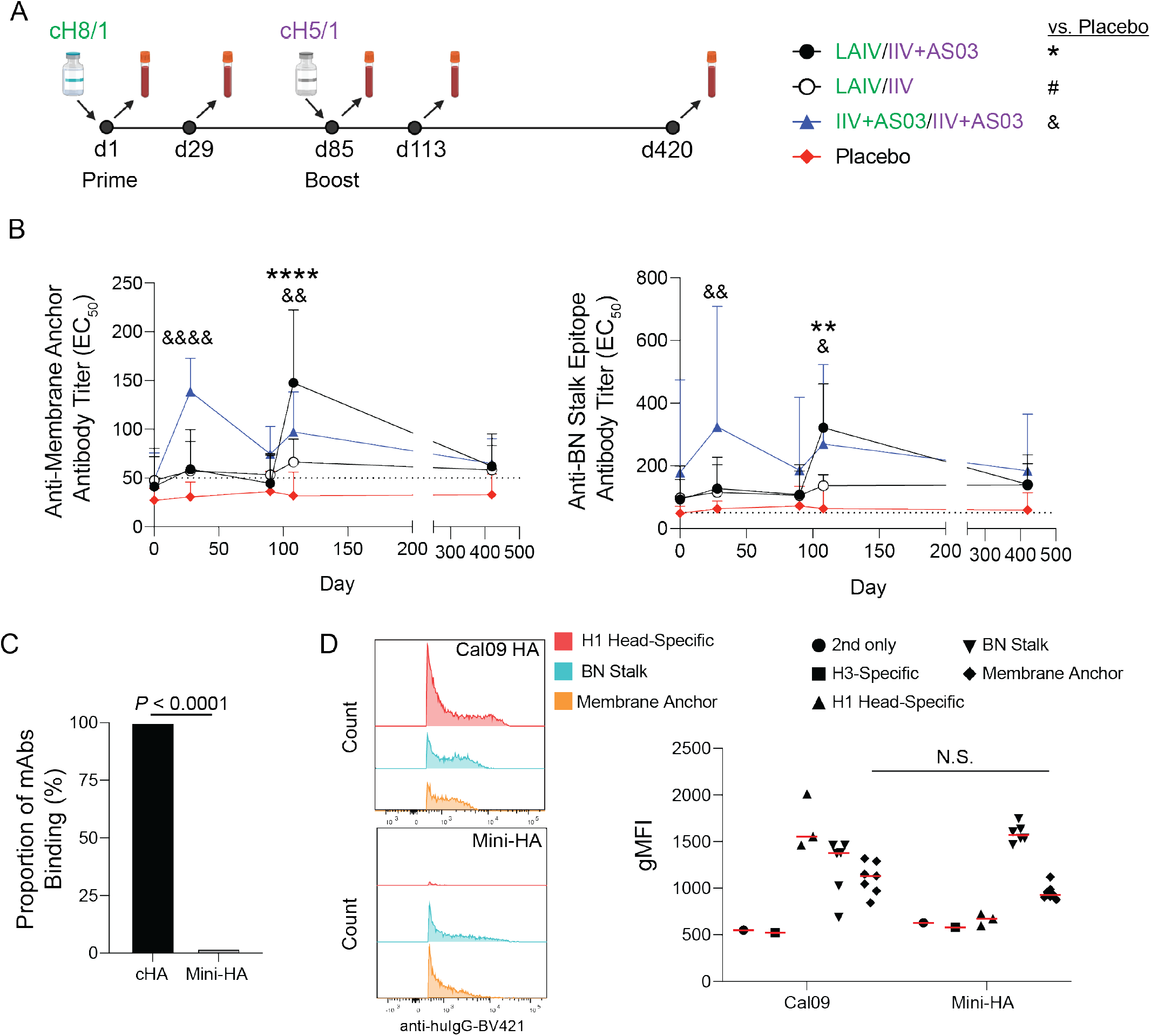
cHA vaccination in humans recalls MBCs targeting the anchor epitope. (**A**-**B**) Subjects enrolled in a phase 1 clinical trial received a prime-boost of cHA vaccine, where the prime used cH8/1 and the boost used cH5/1. On the prime, subjects either received a LAIV or IIV with adjuvant (AS03). On the boost, subjects received the IIV with or without adjuvant (AS03). Serum was collected before and after vaccination and monitored for competing serum antibodies against the anchor epitope (047-09 4F04) and BN stalk epitope (CR9114). LAIV/IIV+AS03 (n=10); LAIV/IIV (n=7); IIV+AS03/IIV+AS03 (n=7); Placebo (n=6). (**A**) Trial design. (**B**) EC50s of serum antibodies competing for binding with 047-09 4F04 for binding to the anchor epitope (left) and CR9114 for binding to the BN stalk epitope (right). Data are mean + S.D. (**C**) Proportion of anchor epitope binding mAbs binding to cHA or mini-HA. (**D**) MAb binding to HEK293T cells expressing full length A/California/7/2009 HA (Cal09) or mini-HA with a transmembrane domain. Representative flow cytometry plots of mAbs binding to Cal09 HA and mini-HA (left) and geometric mean fluorescence intensity (gMFI) of mAbs binding to Cal09 and mini-HA (right). Data represent the median and each symbol represents a distinct mAb. Data in **B** were analyzed using a two-way ANOVA testing for simple effects within rows, data in **C** were analyzed by Fisher’s Exact test and data in **D** were analyzed by unpaired non-parametric Mann-Whitney test. See also **Figure S5.**

Headless HA antigens, or mini-HAs, are attractive universal influenza virus vaccine antigens, as these antigens lack the immunodominant epitopes of the HA head (Impagliazzo et al., 2015; Yassine et al., 2015). We next tested whether the anchor epitope was present on the recombinant mini-HA antigen (Impagliazzo et al., 2015) by performing ELISAs with the anchor epitope targeting antibodies. Only 1 out of 50 anchor antibodies bound the mini-HA antigen, whereas all anchor epitope binding mAbs bound cH6/1 (Figure 5C; Figure S5E). In contrast, all but one BN stalk epitope targeting mAbs were capable of binding to both the mini-HA and the cH6/1 (Figure S5E), suggesting the anchor epitope is specifically disrupted on this antigen. Compared to full-length HA, the membrane proximal region of the mini-HA splays by approximately 14.5 Å (Impagliazzo et al., 2015), suggesting this splaying may disrupt the antigenicity of the anchor epitope. Notably, the mini-HA antigen utilizes a GCN4 trimerization domain, whereas the cH6/1 utilizes a fibritin trimerization domain. Therefore, we next tested whether the loss of antigenicity of the anchor epitope could be due to the utilization of the GCN4 trimerization domain. We identified that anchor epitope targeting antibodies could bind A/California/7/2009 rHA with a fibritin trimerization domain, but not A/California/7/2009 rHA with a GCN4 trimerization domain (Figure S5F), indicating a GCN4 trimerization domain affected the antigenicity of this epitope. To understand whether anchor epitope targeting antibodies could bind the mini-HA in a more native setting, we modified the mini-HA antigen to remove the trimerization domain and include a transmembrane domain, which would lead to the HA being membrane-bound. Transfected HEK293T cells were stained with mAbs targeting H1 head epitopes, the BN stalk epitope, or the anchor epitope, and flow cytometry was performed. MAbs targeting the anchor and BN stalk epitopes readily bound both the full-length membrane-bound A/California/7/2009 HA (Cal09) and the membrane-bound mini-HA, whereas the H1 head-specific mAbs only bound the full-length Cal09 HA (Figure 5D). These data indicate that the anchor epitope is antigenic when HA is trimerized more similarly to membrane-bound HA. Additionally, these data indicate native-like HA antigens are likely to recall MBCs targeting the anchor epitope, such as the cHA vaccine candidate.

### Anchor epitope targeting mAbs utilize a restricted antibody repertoire

We next investigated the repertoire features of mAbs targeting the anchor epitope. All mAbs that targeted the anchor epitope utilized one of four VH3 genes: VH3-23, VH3-30/VH3-30-3, and VH3-48, with over three-quarters utilizing VH3-23 (Figure 6A-B). MAbs targeting the BN stalk epitope commonly used VH1 genes, of which the vast majority used VH1-69 (Figure 6A, C). Anchor epitope targeting mAbs used a variety of DH genes (Table S2) and JH genes, although 70% of anchor targeting mAbs utilized JH4 (Figure S6A). Amongst the anchor targeting mAb heavy chain sequences, 75% were non-clonal (Figure 6D), indicating most anchor targeting utilize similar but distinct heavy chain VDJ recombinations. Similar to the heavy chain, anchor epitope binding mAbs utilized a highly restricted light chain repertoire relative to the BN stalk binding mAbs, with all mAbs utilizing a combination of VK3-11 or VK3-15 combined with JK4 or JK5 (Figure 6E-G; Figure S6B). In contrast, mAbs targeting the BN stalk epitope used a wide array of VK/VL genes (Figure 6E) and JK/JL genes (Figure S6B). Furthermore, all but one light chain of the anchor targeting mAbs were clonal (Figure 6H), indicating the light chains were very similar across mAbs and subjects. By determining paired heavy and light chain clones, we identified 4 distinct clonal expansions, with one public clonal expansion found across multiple subjects (Figure 6I-J; Figure S6C). Anchor epitope targeting mAbs were mutated to a similar extent as mAbs targeting the BN stalk epitope (Figure S6D). The K-CDR3 length of anchor epitope binding mAbs was highly restricted, with all K-CDR3s being ten amino acids in length (Figure S6E). Together, these data indicate that anchor epitope targeting mAbs utilize a highly restricted repertoire, particularly for the light chain.

**Figure 6:**
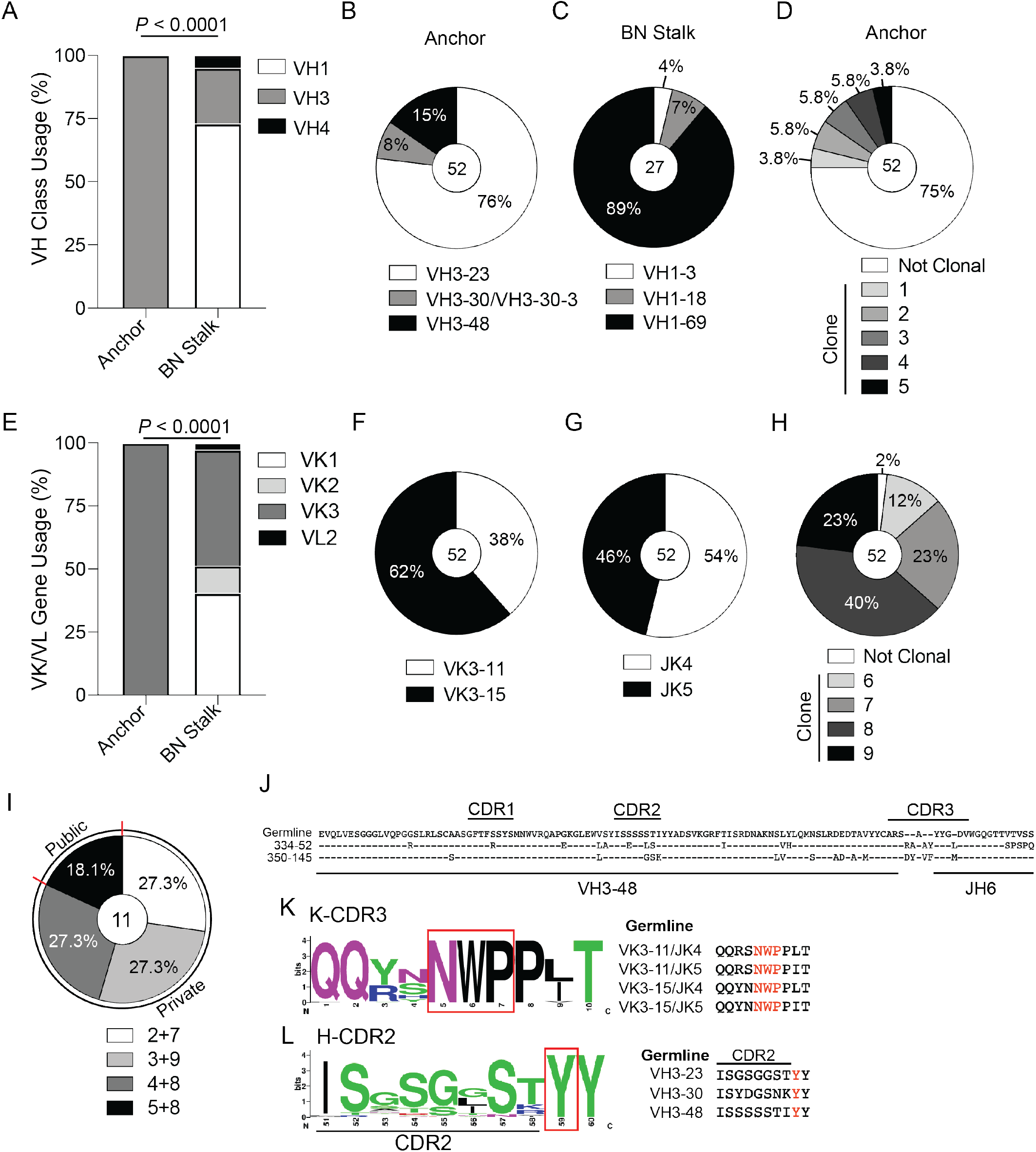
Anchor targeting mAbs use a highly restricted repertoire and possess a conserved binding motif within the K-CDR3. (**A**-**C**) Heavy chain VH classes (**A**) and gene usage of mAbs binding the anchor epitope (**B**) and the BN stalk epitope (**C**). Only VH1 gene usage of BN stalk epitope binding mAbs is shown in **C**. (**D**) Heavy chain clonality of mAbs binding the anchor epitope. (**E**) Light chain VK/VL classes usage of mAbs binding the anchor epitope or BN stalk epitope. (**F**-**G**) VK (**F**) and JK (**G**) gene usage of mAbs binding the anchor epitope. (**H**) Light chain clonality of mAbs binding the anchor epitope. (**I**) Private and public clones that share heavy and light chains. (**J**) Alignment of VDJ of the VH3-48 public clone. (**K**) Sequence logo of the K-CDR3 and the germline sequence of the K-CDR3 of VK3-11/VK3-15 combined with JK4/JK5. NWP motif is highlighted. (**L**) Sequence logo of the H-CDR2 with the tyrosines directly following the H-CDR2 and the germline sequence of the H-CDR2 of VH3-23, VH3-30, and VH3-48. Data in **A** and **E** were analyzed using Chi-square tests. See also **Figure S6**.

FISW84 (Benton et al., 2018) similarly uses VH3-23/VK3-15 and largely makes interactions with the epitope via an NWP motif within the K-CDR3 loop and a tyrosine (Y) immediately following the H-CDR2 (Figure S6F). We identified that all anchor targeting mAbs possessed this NWP motif at the exact same location within the K-CDR3, which was present in the germline sequence of the various VK/JK pairings (Figure 6K). Moreover, all anchor binding mAbs utilized a germline encoded tyrosine at position 59 (Figure 6L), suggesting this residue could have led to the selection of B cells utilizing these particular VH3 genes. Despite this, nearly 2/3 of VH3 genes utilize a tyrosine at this exact position (Figure S6G), suggesting other features of the heavy chain may lead to the preferential selection of these particular VH3 genes into the B cell repertoire against the anchor epitope. Together, these data reveal B cells targeting the anchor epitope utilized a highly restricted V(D)J gene repertoire, and these specific features of the repertoire are likely critical for binding the anchor epitope.

### Humans possess MBCs with features of anchor epitope targeting antibodies

Due to the restricted repertoire features of anchor targeting mAbs, we next determined the relative proportion of B cell subsets with these features by integrating single-cell RNA-sequencing and repertoire sequencing of HA-specific B cells isolated from 22 subjects following cH5/1 vaccination (d112; Figure 5A). Notably, most B cells isolated likely target the H1 stalk domain as we sequenced sorted cH5/1^+^ B cells and humans have no measurable pre-existing immunity against the H5 head domain (Han et al., 2020). However, subjects may have recruited naïve B cells against the H5 head component of the cH5/1 vaccine, therefore the isolated B cell pool is likely a heterogenous population of mostly H1 stalk domain-reactive B cells and some H5 head domain-reactive B cells. To investigate the proportion of B cells with repertoire features of B cells targeting the anchor epitope, we selected B cells that used VH3-23/VH3-30/VH3-30-3/VH3-48, VK3-11/VK3-15, JK4/JK5, a 10 amino acid length K-CDR3, and possessed an NWP motif within the K-CDR3. For reference, we additionally segregated out B cells expressing VH1-69 and a kappa chain, as these are the dominate repertoire features of B cells targeting the BN stalk epitope (Figure 6A, C, E). We identified that B cells with features of antibodies binding the anchor epitope were abundant within the human B cell repertoire, with 6% of all B cells identified fitting within this defined repertoire (Figure 7A). Of subjects with ten or more VDJ^+^ B cells (n=20), we identified anchor targeting B cells in all but one subject (Figure 7B). The anchor targeting B cell pool largely used VH3-23/VK3-15 pairing (Figure 7C). Additionally, we generated 34 mAbs from the selected anchor targeting B cell list and 31 of these mAbs competed with 047-09 4F04 (Figure 7D). The anchor epitope B cells had a similar number of mutations as VH1-69/kappa B cells (Figure 7E) and were largely class-switched to IgG1 and IgG3 (Figure 7F), indicative of prior class-switch recombination and B cell selection within germinal centers. Together, these data indicate that the anchor epitope is a common target of the human MBC repertoire against HA. Moreover, this study indicates that most adults have pre-existing immunity against this epitope that can be harnessed by a potential universal influenza virus vaccine candidate to provide broad protection against H1-expressing viruses.

**Figure 7:**
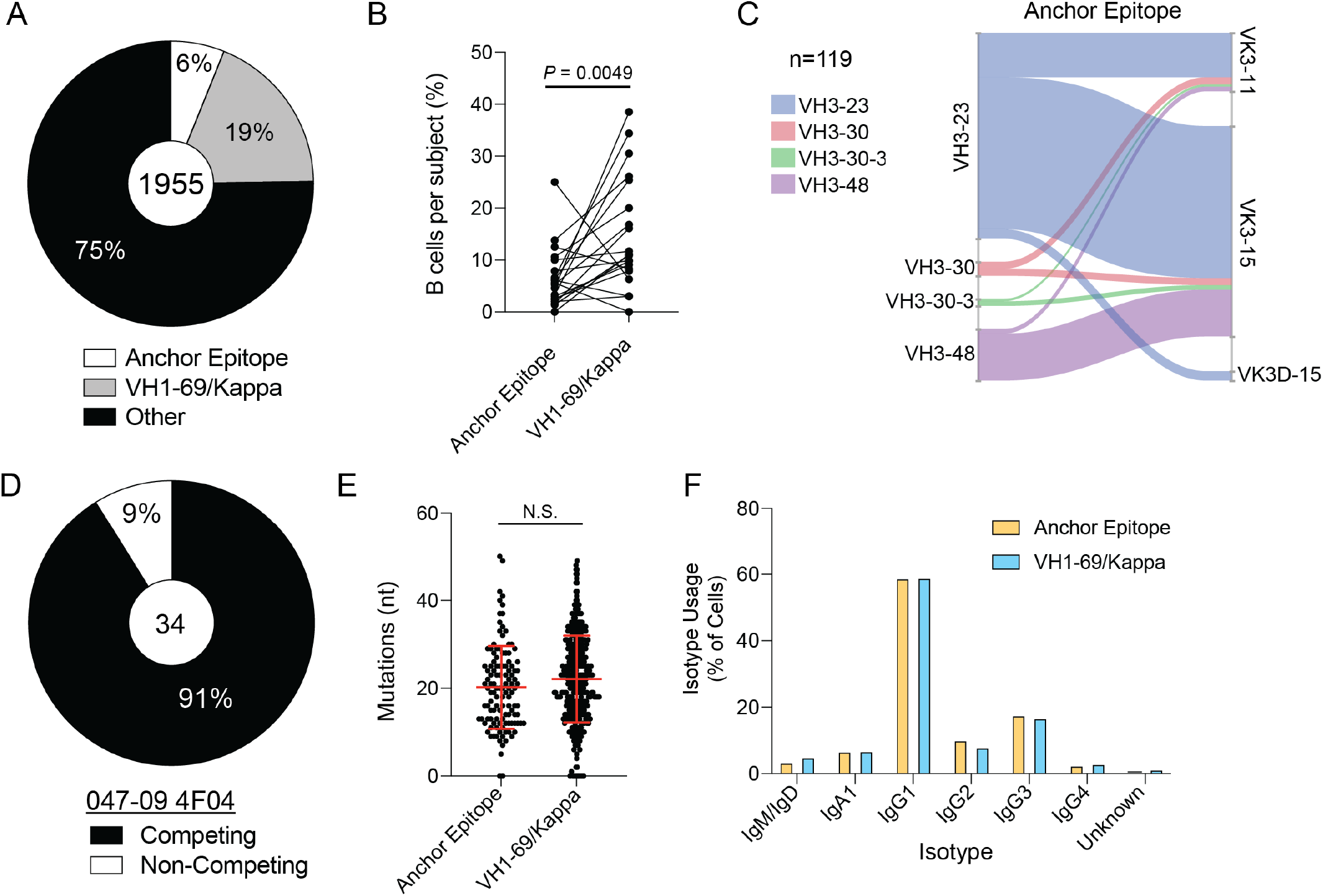
Humans possess MBCs targeting the anchor epitope. cH5/1^+^ B cells from PBMCs were sorted from subjects 28 days following a booster with the cH5/1 and were subjected to single-cell RNA-sequencing. (**A**) Proportion of all B cells with features of anchor antibodies, VH1-69/kappa (BN stalk epitope), or with other repertoire features. (**B**) Proportion of B cells with anchor binding antibody features or that use VH1-69/kappa chain by subject. Lines connect the same subject. (**C**) VH/VK pairing of B cells with features of anchor epitope binding antibodies. (**D**) 34 mAbs with anchor epitope binding mAb repertoire features and tested for competing for binding with 047-09 4F04. (**E**-**F**) number of heavy chain mutations (**E**) and isotype usage (**F**) of B cells with repertoire features of anchor binding antibodies or VH1-69/kappa. Data in **E** are represented as mean ± S.D. Data in **D** were analyzed using a paired non-parametric Wilcoxon matched-pairs signed rank test and data in **E** were analyzed by unpaired non-parametric Mann-Whitney test.

## Discussion

In this study, we identified a class of antibodies targeting a broadly neutralizing epitope of hemagglutinin stalk domain near the viral membrane of H1-expressing influenza viruses. The stalk domain is conserved within and often across influenza virus subtypes. Anchor epitope targeting antibodies showed broad neutralizing activity against H1-expressing influenza viruses but rarely cross-reacted with other influenza subtypes. The anchor epitope was poorly conserved across influenza virus subtypes, which could explain the H1 subtype specificity of the anchor epitope targeting antibody class identified in this study. However, the broadly neutralizing activity of anchor epitope targeting mAbs against pre- and post-pH1N1 viruses and a swine-origin H1-expressing virus indicates the anchor epitope is an important target for pan-subtype neutralizing antibodies. Anchor epitope targeting antibodies have the potential to provide protection against antigenically drifted H1N1 viruses and zoonotic spillovers of H1-expressing viruses. Furthermore, stalk binding antibodies are an independent correlate of protection against influenza virus infection and lower respiratory symptoms (Aydillo et al., 2020; Ng et al., 2019). Whether antibodies against distinct stalk domain epitopes are independent correlates of protection against influenza virus infection is yet to be determined.

A striking feature of anchor epitope binding antibodies was the angle of approach, with the antibody Fab tilting up towards the epitope and sterically clashing with the viral membrane. However, our data indicate anchor epitope antibodies can bind HA in the context of the viral membrane, as these antibodies could bind intact virus and were neutralizing *in vitro*. These data are consistent with a dynamic HA on the membrane, tilting up to 52° along its three-fold axis (Benton et al., 2018), allowing for the exposure of epitopes proximal to the membrane. As HA flexes on the viral membrane surface, the epitope may become available, allowing for antibody binding at the observed angle. However, likely only 1-2 of the anchor epitopes of the HA trimer is accessible as HA flexes, limiting the avidity of antibodies binding the anchor epitope. Moreover, influenza viruses are densely decorated with the surface glycoproteins HA and NA (Gallagher et al., 2018; Wasilewski et al., 2012), further limiting access to the anchor epitope. Further research is needed to understand how the anchor epitope can be made more accessible for antibody binding. Moreover, it is of interest to understand whether anchor epitope binding antibodies possess membrane binding capabilities, similar to antibodies binding to membrane proximal external region (MPER) of gp41 of HIV (Cardoso et al., 2005; Ofek et al., 2004).

The anchor targeting mAbs utilized a highly restricted repertoire that were public clonotypes across subjects, with all antibodies possessing two conserved motifs near the H-CDR2 and a NWP motif within the K-CDR3. In addition, VH3-23 and VH3-48 utilizing mAbs targeted the anchor epitope slightly differently, with the VH3-48 mAb (047-09 4F04) sitting more anterior and superior on a single HA protomer relative to the VH3-23 mAb (241 IgA 2F04). However, more antibodies need to be studied to further understand how the slight differences in repertoire usage affect HA binding. Multiple classes of antibodies against the BN stalk epitope of H1-expressing viruses have been identified (Joyce et al., 2016; Sui et al., 2009) and often cross-react with other group 1 virus subtypes (i.e. H2, H5), and occasionally group 2 viruses (Henry Dunand et al., 2015). Notably, each antibody class targets the BN stalk epitope at slightly different angles and have slightly different binding footprints (Joyce et al., 2016; Wu and Wilson, 2020). As a result of this, viral mutations that arise to circumvent one antibody class may have a minimal effect on viral binding breadth and neutralization potential of other classes. Moreover, naturally occurring mutations within the footprint of the anchor antibodies have not been observed, whereas mutations against the BN stalk epitope have been observed (Wu et al., 2020).

Our study showed that humans have pre-existing immunity against the anchor epitope and influenza virus vaccination can recall MBCs to secrete antibodies against this epitope. However, vaccine HA antigens must have a native confirmation near the transmembrane domain, as our study showed that trimer splaying potentially due to the GCN4 trimerization domain ablates antibody binding at the anchor epitope. Split vaccines, HA-decorated nanoparticles, and mRNA vaccines that induce expression of membrane-bound HA should possess a native anchor epitope that can be recognized by MBCs targeting this epitope. Moreover, our study highlights that the cHA vaccine strategy was able to recall MBCs against the anchor epitope and the BN stalk epitope, while avoiding the recruitment of MBCs targeting the variable epitopes of the HA head (Nachbagauer et al., 2020). Similarly, the mini-HA/headless HA vaccine strategy has the potential to also recall MBCs against multiple epitopes of the HA stalk domain, if folded natively (van der Lubbe et al., 2018; Yassine et al., 2015). However, an optimal pan-H1 vaccine should strive to induce antibodies against both conserved epitopes of the HA stalk domain and head domain, including the RBS, lateral patch, and trimer interface in order to limit potential viral escape mutants. Vaccines that strategically glycosylate variable epitopes have the potential to induce antibodies against conserved epitopes of the HA head and stalk domains, while limiting B cell recruitment against variable epitopes (Bajic et al., 2019; Boyoglu-Barnum et al., 2020; Eggink et al., 2014; Weidenbacher and Kim, 2019). Moreover, mosaic antigens that replace the variable epitopes with those from avian influenza virus subtypes also have the potential to induce broadly protective antibodies against HA (Broecker et al., 2019; Liu et al., 2018; Sun et al., 2019). Together, our study indicates that novel influenza vaccination strategies have the capability to robustly induce antibodies against the previously unappreciated anchor epitope that can provide broad protection against H1-expressing viruses.

## Supporting information

Supplement Figures, Tables, and References

## Acknowledgments

We are thankful to all subjects who participated in this study. We thank Sarah Andrews, Rafi Ahmed, Jens Wrammert, and Karlynn Neu for initiating studies on the 2009 MIV, 2010 TIV, and 2014 QIV cohorts. We thank Chiara Mariottini, Jodi Feser, Daniel Stadlbauer, and Anna-Karin Palm for their help on the cHA vaccine trial. We thank Ian Wilson and Alec Freyn for fruitful discussion and feedback on experimental design. We are thankful to the teams at PATH, GSK, Cincinnati Children’s Hospital Medical Center, and Duke University for their work on the chimeric HA vaccine trial (NCT03300050), which was funded in part by the Bill and Melinda Gates Foundation (OPP1084518). The findings and conclusions contained within are those of the authors and do not necessarily reflect positions or policies of the Gates Foundation. This project was funded in part by the National Institute of Allergy and Infectious Diseases; National Institutes of Health grant numbers U19AI082724 (P.C.W.), U19AI109946 (P.C.W.), U19AI057266 (P.C.W.), P01 AI097092 (P.P.), R01AI145870-01 (P.P.), R21AI146529 (L.C), and T32AI007244-36 (J.H.), and the NIAID Centers of Excellence for Influenza Research and Surveillance (CEIRS) grant number HHSN272201400005C (P.C.W.), and HHSN272201400008C (L.C., F.K., A.G.-S., P.P.). This work was also partially supported by the National Institute of Allergy and Infectious Disease (NIAID) Collaborative Influenza Vaccine Innovation Centers (CIVIC; 75N93019C00051, F.K., A.G.-S., P.P., A.B.W., P.C.W.).

## Author contributions

J.J.G. designed the study, characterized mAbs, analyzed the data, and wrote the manuscript. J.H. generated structures, performed mapping experiments, analyzed data, and edited the manuscript. H.A.U. performed characterization ELISAs. L.L. analyzed single-cell RNA-sequencing data. L.Y.L. sorted cH5/1^+^ B cells and generated RNA-sequencing data. C.H. and C.T.S. generated mAbs from cHA vaccine trial. O.S., D.J.B., and S.C. performed virus-specific ELISAs. J.J.G, H.L.D., M.E.T., C.T.S., and H.A.U. performed infection challenge studies. L.G. and J.D.B. generated HA mutant data on 045-09 2B06. N.-Y.Z. grew and purified influenza viruses. S.T.R. helped perform EMPEM studies. M.H. performed mAb cloning. S.S. and F.K. provided recombinant proteins. F.K., P.P., A.G.-S., and R.N. designed and orchestrated cHA vaccination trial. L.C. provided recombinant protein and plasmids for membrane-bound HA. A.B.W. supervised structural analyses and provided critical feedback on experimental design, P.C.W. supervised the work and edited the manuscript. All authors provided feedback on the manuscript.

## STAR METHODS

### KEY RESOURCES TABLE

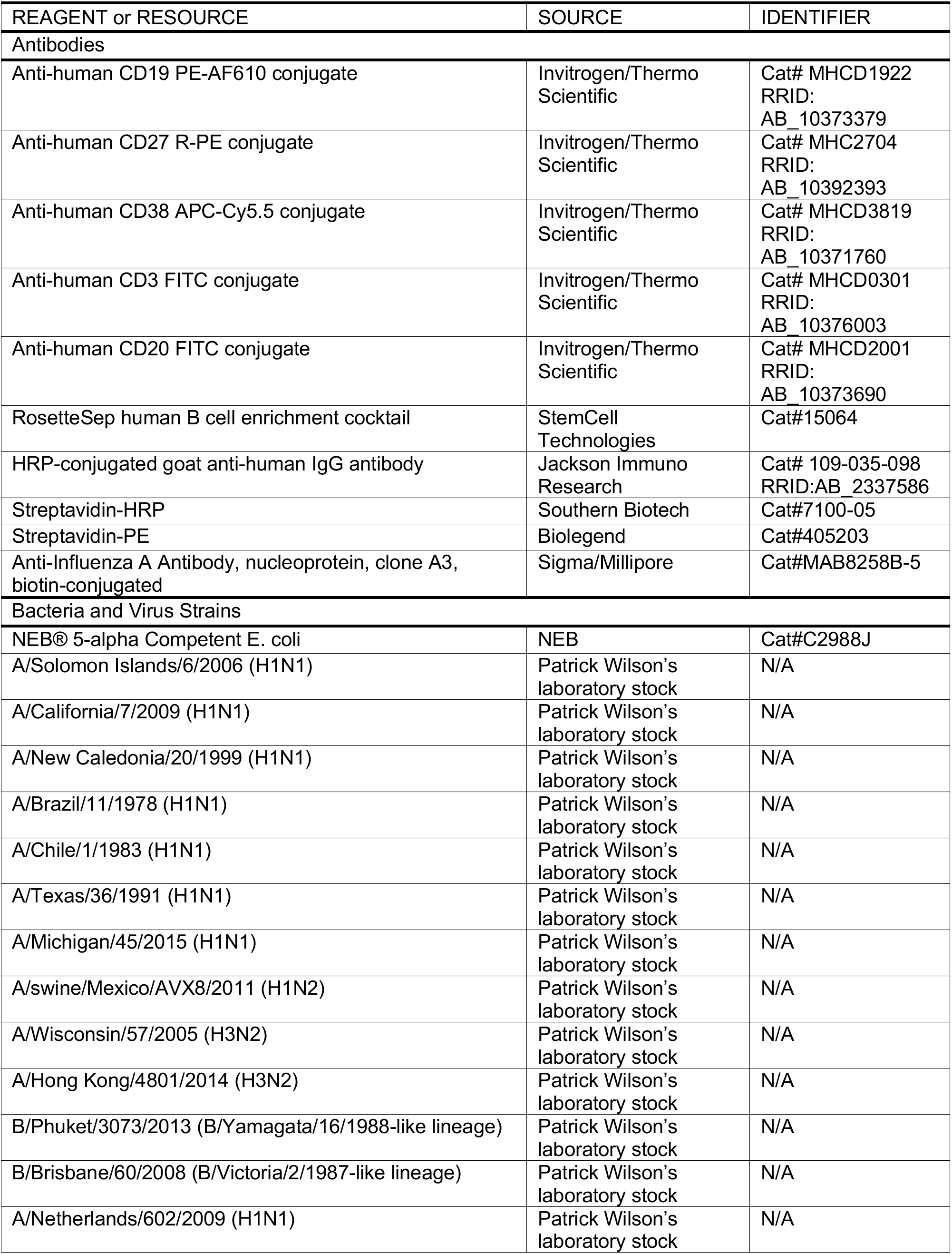
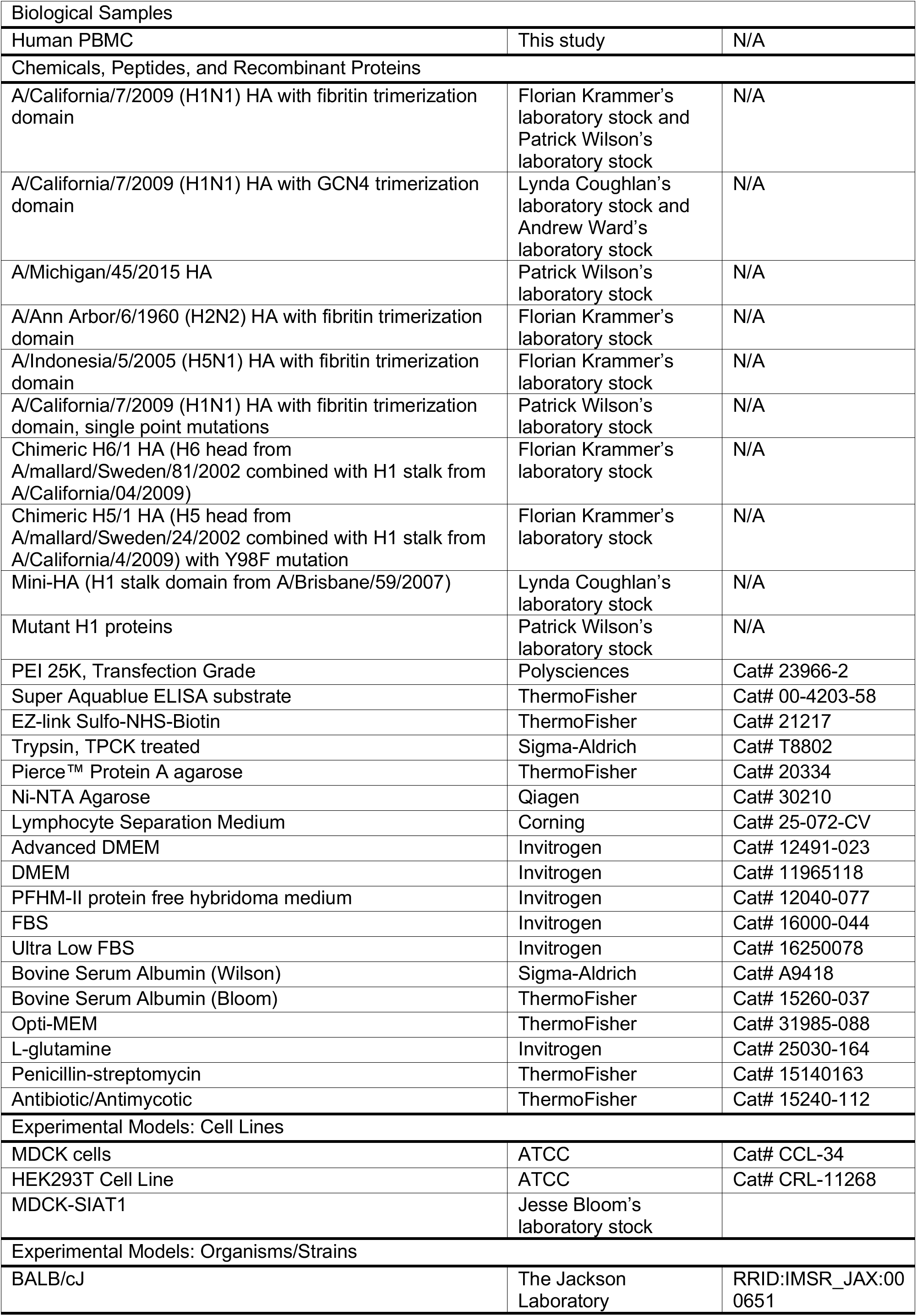
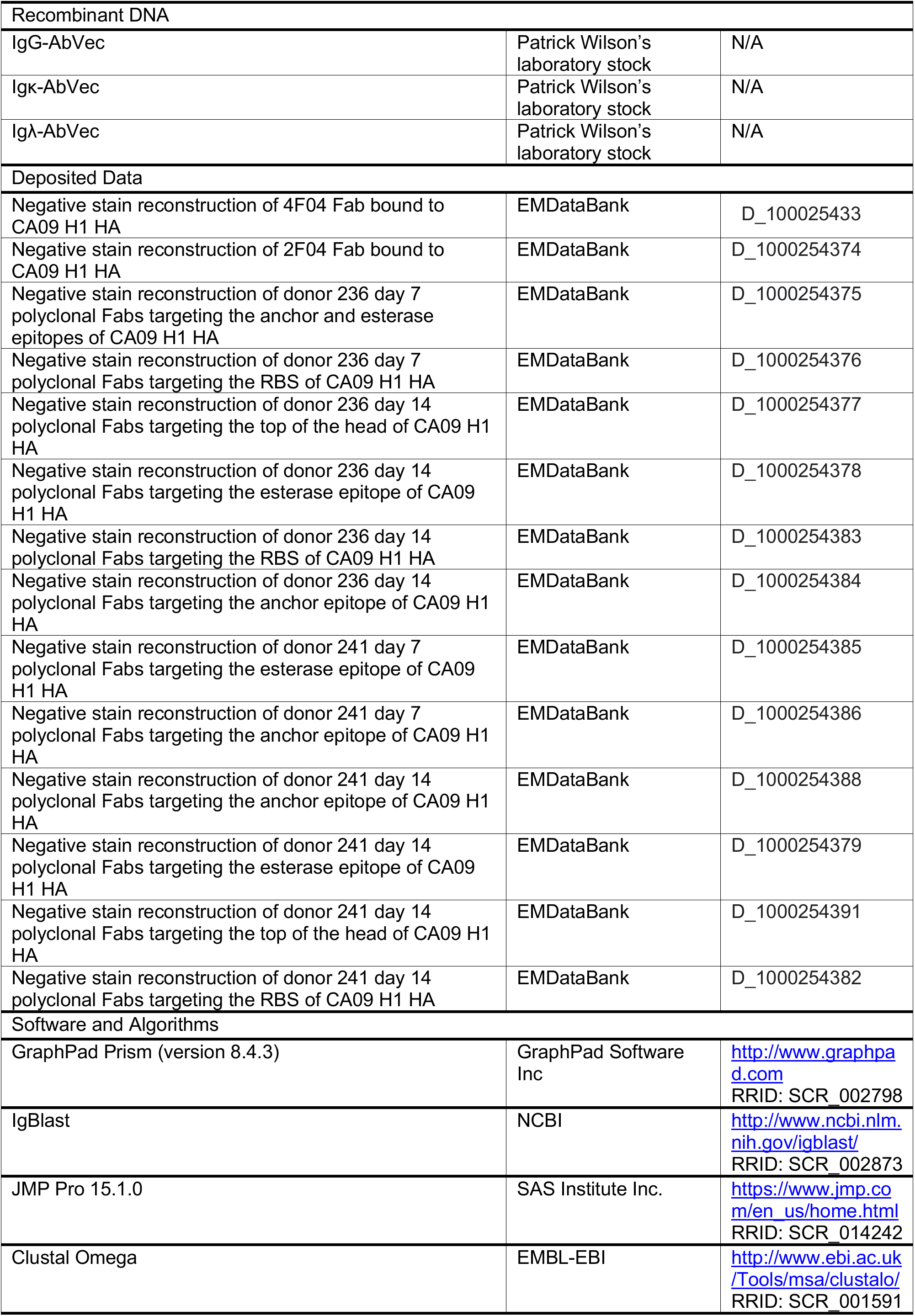
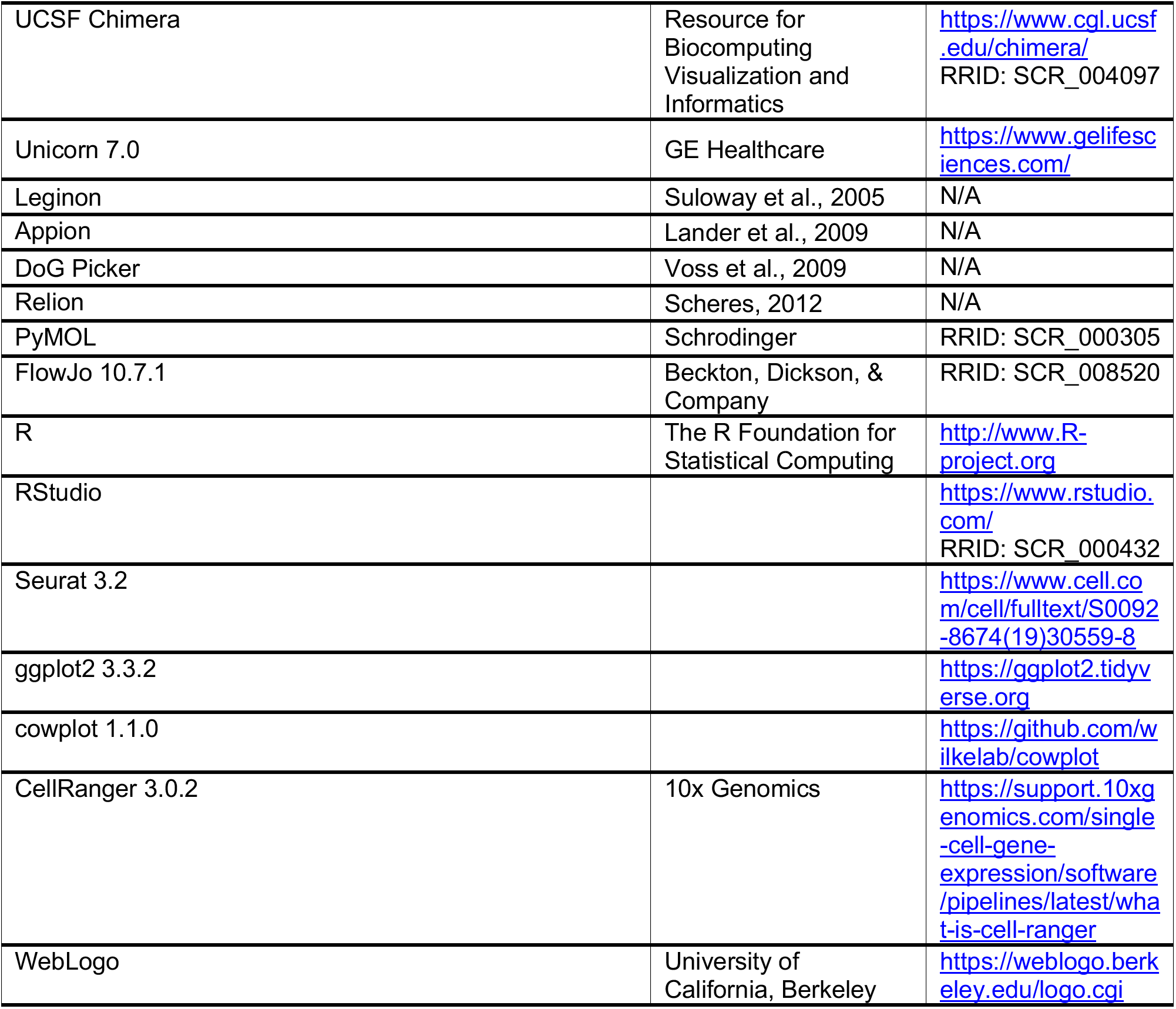

#### Resource Availability

##### Lead Contact

Further information and requests for resources and reagents should be directed to the Lead contact, Patrick C. Wilson.

##### Materials Availability

There are restrictions to the availability of mAbs from this study due to the lack of an external centralized repository for its distribution and our need to maintain the stock. We are glad to share mAbs with reasonable compensation by requestor for its processing and shipping.

##### Data and Code Availability

Repertoire data generated from single cell RNA-sequencing data is deposited at NCBI GenBank under accession numbers (in process of being deposited). Electron microscopy maps were deposited to the Electron Microscopy DataBank under accession IDs: D_100025433, D_1000254374, D_1000254375, D_1000254376, D_1000254377, D_1000254378, D_1000254383, D_1000254384, D_1000254385, D_1000254386, D_1000254388, D_1000254379, D_1000254391, and D_1000254382. All next generation sequencing data for 045-09 2B06 deep mutational scanning can be found on the Sequence Read Archive under BioProject accession number PRJNA494885.

### EXPERIMENTAL MODEL AND SUBJECT DETAILS

#### Human Materials

Human PBMCs were obtained from multiple subjects from multiple cohorts, which is outlined in Table S1. All studies were performed with the approval of the University of Chicago Institutional Review Board (ID #09-043-A). The chimeric HA vaccine study cohort is identified as clinical trial NCT03300050.

#### Cell Lines

Human Embryonic Kidney HEK293T (female, # CRL-11268) and Madin Darby canine kidney MDCK (female, # CCL-34, NBL-2) cells were purchased and authenticated by the American Type Culture Collection (ATCC). All cells were maintained in a humidified atmosphere of 5% CO2 at 37°C. HEK293T cells were maintained in Advanced-DMEM supplemented with 2% ultra-low IgG fetal bovine serum (FBS) (Invitrogen), 1% L-glutamine (Invitrogen) and 1% antibiotic-antimycotic (Invitrogen). MDCK cells were maintained in DMEM supplemented with 10% FBS (Invitrogen), 1% L-glutamine (Invitrogen) and 1% penicillin-streptomycin (Invitrogen).

### METHOD DETAILS

#### Monoclonal antibody production

Monoclonal antibodies were generated as previously described (Guthmiller et al., 2019; Smith et al., 2009; Wrammert et al., 2008). Peripheral blood was obtained from each subject approximately 7 days after vaccination or infection or obtained 28+ days post-vaccination. Lymphocytes were isolated and enriched for B cells using RosetteSep. Total PBs (CD3^−^CD19^+^CD27^hi^CD38^hi^; all cohorts except 2014 QIV), IgG^+^ PBs (CD3^−^CD19^+^IgM^−^CD27^hi^CD38^hi^IgG^+^IgA^−^; 2014 QIV), IgA^+^ PBs (CD3^−^CD19^+^IgM^−^CD27^hi^CD38^hi^IgG^−^IgA^+^; 2014 QIV cohort), or HA^+^ bait-sorted MBCs (CD3^−^CD19^+^CD27^+^CD38^lo/+^HA^+^, for 030-09M 1B06) were single-cell sorted into 96-well plates. Immunoglobulin heavy and light chain genes were amplified by reverse transcriptase polymerase chain reaction (RT-PCR), sequenced, cloned into human IgG1, human kappa chain, or human lambda expression vectors, and co-transfected into HEK293T cells. Secreted mAbs were purified from the supernatant using protein A agarose beads. For mAbs generated from the 2014 QIV cohort, mAb names include the original isotype of the sorted PB, and all mAbs were expressed as human IgG1. cH5/1-binding B cells (CD19^+^CD27^+^cH5/1^+^) were sorted from subjects 28 days after cH5/1 vaccination (NCT03300050). Cells were sorted with A/California/04/2009 HA (for 030-09M 1B06) or cH5/1 probe with a Y98F mutation to ablate non-specific binding to sialic acids on B cells. MAb heavy chain and light chain sequences were synthesized from single-cell RNA-sequencing data of cH5/1-baited B cells (IDT), and cloned into the human IgG1, human kappa chain, or human lambda expression vectors. B cell clones were determined by aligning all the V(D)J sequences sharing identical progenitor sequences, as predicted by IgBLAST using our in-house software, VGenes. Consensus sequence analysis was performed using WebLogo (Crooks et al., 2004) and sequence alignments were determined using Clustal Omega.

#### Viruses and recombinant proteins

Influenza viruses used in all assays were grown in-house in specific pathogen free (SPF) eggs, harvested, purified, and titered. The A/swine/Mexico/AVX8/2011 H1N2 virus (Mena et al., 2016) was provided by Ignacio Mena, Adolfo García-Sastre, and Sean Liu at Icahn School of Medicine at Mount Sinai. Recombinant HA, cHA, and mini-HA were obtained from BEI Resources or kindly provided by the Krammer laboratory at Icahn School of Medicine at Mount Sinai, the Coughlan laboratory at The University of Maryland School of Medicine, or the Wilson laboratory at the University of Chicago. Recombinant HA mutant proteins used in Figure 2 were generated with identified mutations from the deep mutational scanning experiments (see below) or with known mutations that have arisen naturally or were identified in other studies (Figure S3). All mutations were made on HA from A/California/7/2009. Mutant HAs were expressed in HEK293T cells and purified using Ni-NTA agarose beads (Qiagen).

#### Antigen Specific ELISA

High protein-binding microtiter plates (Costar) were coated with 8 hemagglutination units (HAU) of virus in carbonate buffer or with recombinant HA, including HA mutants described below, at 2 μg/ml in phosphate-buffered saline (PBS) overnight at 4°C. Plates were washed the next morning with PBS 0.05% Tween and blocked with PBS containing 20% fetal bovine serum (FBS) for 1 hr at 37°C. Antibodies were then serially diluted 1:3 starting at 10 μg/ml and incubated for 1.5 hr at 37°C. Horseradish peroxidase (HRP)-conjugated goat anti-human IgG antibody diluted 1:1000 (Jackson Immuno Research) was used to detect binding of mAbs, and plates were subsequently developed with Super Aquablue ELISA substrate (eBiosciences). Absorbance was measured at 405 nm on a microplate spectrophotometer (BioRad). To standardize the assays, control antibodies with known binding characteristics were included on each plate, and the plates were developed when the absorbance of the control reached 3.0 OD units. All ELISAs were performed in duplicate twice. To determine mAb affinity, a non-linear regression was performed on background subtracted ODs and Kd values were reported. To classify antigen-specificity, mAbs that did not definitively bind the HA head or stalk are listed as binding unknown HA^+^ epitopes. Affinity measurements, as represented as Kd at a molar concentration (M), were calculated using Prism 9 (Graphpad) by performing a non-linear regression.

#### Deep mutational scanning for stalk domain mutants

The mutant libraires used herein were previously described (Doud and Bloom, 2016). The libraries consist of all single amino-acid mutations to A/WSN/1933 (H1N1). The experiments were performed by using biological triplicate libraries. The mutational antigenic profiling of the 045-09 2B06 was performed as previously outlined (Doud et al., 2017). In brief, 10^6^ TCID_50_ of two of the virus library biological replicates was diluted in 1mL in IGM (Opti-MEM supplemented with 0.01% FBS, 0.3% BSA, and 100 μg/ml calcium chloride) and incubated with an equal volume of 045-09 2B06 antibody at a final concentration of 50 or 25 μg/mL for 1.5 hours at 37°C. MDCK-SIAT1 cells were infected with the virus antibody mixtures. 2 hours post-infection, the media was removed, the cells washed with 1 ml PBS, and 2 ml of fresh IGM was added. 15 hours post-infection, viral RNA was extracted, reverse-transcribed using primers WSNHA-For (5’-AGCAAAAGCAGGGGAAAATAAAAACAAC-3’) and WSNHA-Rev (5’-AGTAGAAACAAGGGTGTTTTTCCTTATATTTCTG-3’), and PCR amplified according to the barcoded-subamplicon library preparation as previously described (Doud and Bloom, 2016). The overall fraction of virions that survive antibody neutralization was estimated using qRT-PCR targeting the viral nucleoprotein (NP) and cellular GAPDH as previously described (Doud et al., 2017). Using 10-fold serial dilutions of the virus libraries, we infected cells with no antibody selection to serve as a standard curve of infectivity. qPCR Ct values from the standard curve samples compared to the virus-antibody mix samples are determined for NP and GAPDH. We then generate a linear regression to fit the difference between the NP and GAPDH Ct values for the standard curve samples, and then use this curve to interpolate the fraction surviving for the antibody-virus selection samples. Across the three library replicates the fraction of virus surviving antibody selection was 0.17, 0.1, and 0.14.

Illumina(R) deep sequencing data was analyzed using dms_tools2 version 2.4.12 software package (Bloom, 2015) which can be found at https://github.com/jbloomlab/dms_tools2. All of the computer code used is at https://github.com/jbloomlab/2B06_DMS, and the Jupyter notebook that performs most of the analysis is at https://github.com/jbloomlab/2B06_DMS/blob/master/analysis_notebook.ipynb. The sequencing counts were processed to estimate the differential selection for each mutation, which is the log enrichment of that mutation in the antibody-selected condition versus the control (Doud et al., 2017). The numerical measurements of the differential selection that 2B06 imposes on each mutation can be found here: https://github.com/jbloomlab/2B06_DMS/blob/master/results/diffsel/tidy_diffsel.csv.

#### Competition ELISAs

Plates were coated with 50μl of A/California/7/2009 HA at a concentration of 1μg/ml and incubated overnight at 4°C. To biotinylate the antibodies with known epitope specificities, CR9114 and 047-09-4F04, were incubated at 4°C with EZ-Link™ Sulfo-NHS-Biotin (Thermo Scientific) for 24h or 48h prior to use, respectively. After blocking the plates with PBS 20% FBS for 1h at 37°C, serum samples were incubated (starting dilution of 1:50 for human serum or 20 μg/ml for mAbs) in the coated wells for 2h at room temperature. Either biotinylated CR9114 or 047-09-4F04 was then added at a concentration equal to twice its Kd and incubated in the wells with the serum or mAbs for 2h at room temperature. The biotinylated antibodies were desalted before addition to remove free biotin using Zeba™ spin desalting columns, 7k MWCO (Thermo Scientific). After washing the plates, wells were incubated with HRP-conjugated streptavidin (Southern Biotech) at 37°C for 1h for detection of the biotinylated antibody. Super Aquablue ELISA substrate (eBiosciences) was then added and absorbance was measured at 405nm on a microplate spectrophotometer (Bio-Rad). To standardize the assays, biotinylated CR9114 or TS-09-4F04 was incubated in designated wells on each plate without any competing serum or mAb, and data were recorded when the absorbance of these wells reached an optical density (OD) of 1 to 1.5 units. After subtracting background, percent competition by serum samples was then determined by dividing a sample’s observed OD by the OD reached by the positive control, subtracting this value from 1, and multiplying by 100. For the serum data, ODs were log transformed and analyzed by non-linear regression to determine EC50 values using Prism software (Graphpad). For Figure 5 and Figure S4, only subjects with serum for all timepoints were included.

#### Microneutralization Assays

Microneutralization assays for mAb characterization were carried out as previously described (Chen et al., 2018; Henry Dunand et al., 2015). MDCK cells were maintained in DMEM supplemented with 10% FBS, 1% penicillin-streptomycin, and 1% L-glutamine at 37°C with 5% CO2. The day before the experiment, 25,000 MDCK cells were added to each well of a 96-well plate. Serial two-fold dilutions of mAb were mixed with an equal volume of 100 TCID50 of virus for 1 hr and added to MDCK cells for 1 hr at 37°C. The mixture was removed, and cells were cultured for 20 hrs at 37°C with 1X MEM supplemented with 1 μg/ml tosyl phenylalanyl chloromethyl ketone (TPCK)-treated trypsin and appropriate mAb concentration. Cells were washed twice with PBS, fixed with 80% ice cold acetone at 20°C for at least 1 hr, washed 3 times with PBS, blocked for 30 min with 3% BSA, and then treated for 30 min with 2% H2O2. Cells were incubated with a mouse anti-nucleoprotein antibody (1:1000; Millipore) in 3% BSA-PBS for 1 hr at room temperature (RT), followed by goat anti-mouse IgG HRP (1:1000; Southern Biotech) in 3% BSA-PBS for 1 hr at RT. The plates were developed with Super Aquablue ELISA substrate at 405 nm until virus only controls reached an OD of 1. The signal from uninfected wells was averaged to represent 100% inhibition. The signal from infected wells without mAb was averaged to represent 0% inhibition. Duplication wells were used to calculate the mean and SD of neutralization, and inhibitory concentration 50 (IC50) was determined by a sigmoidal dose response curve. The inhibition ratio (%) was calculated as below: ((OD Pos. Control – OD Sample) / (OD Pos. Control – OD Neg. Control)) * 100. The final IC50 was determined using Prism software (GraphPad).

#### *In vivo* challenge infections

MAb cocktails (Table S4) were passively transferred into 6- to 8-week-old female BALB/c mice (Jackson Laboratories) by intraperitoneal injection of 0.2, 1, and 5 mg/kg mAb cocktail, which are further detailed in Figure S4. Negative control mice received 5 mg/kg of the anthrax-specific mAb 003-15D03 as an isotype control. Two hours post-mAb injection, mice were anesthetized with isoflurane and intranasally challenged with 10 LD50 of mouse-adapted A/Netherlands/602/2009 H1N1 virus, with 10 μl of virus administered into each nostril (20 μl total). As a read out, survival and weight loss were monitored 1-2 times daily for two weeks. Mice were euthanized upon 25% weight loss or at the end of the experiment (14 days post challenge). All experiments were done in accordance with the University of Chicago Institutional Animal Care and Use Committee.

#### HA footprint mapping

The footprints of three mAbs (FISW84 (PDB: 6HJQ), CR9114 (PDB: 4FQI), and FI6v3 (PDB: 3ZTN)) were mapped onto one HA protomer (A/California/4/2009, PDB: 4M4Y) using UCSF Chimera (Pettersen et al., 2004) and Adobe Photoshop. EM maps of HA:fab complexes were aligned in UCSF Chimera and footprints were mapped onto one HA protomer. Individual protomers of the HA trimer are indicated in different shades of gray.

#### Negative stain EM

Immune complexes were prepared by incubating Fab with HA (A/California/04/2009 with E47K or E47G stabilizing mutations) at greater than 3:1 molar ratio for 2 hours at room temperature (RT). Samples were deposited at ~10μg/mL on glow-discharged, carbon-coated 400 mesh copper grids (Electron Microscopy Sciences, EMS) and stained with 2% w/v uranyl formate. Samples were imaged at 52,000x magnification, 120kV, on a Tecnai Spirit T12 microscope equipped with an Eagle CCD 4k camera (FEI) or 62,000 magnification, 200kV, on a Tecnai T20 microscope equipped with a CMOS 4k camera (TVIPS). Micrographs were collected with Leginon, single particles were processed with Appion and Relion, footprints were mapped with UCSF Chimera, and figures were made with UCSF Chimera (Lander et al., 2009; Pettersen et al., 2004; Scheres, 2012; Suloway et al., 2005).

### EMPEM

Human serum samples were heat-inactivated at 55°C for 30min before incubating on Capture Select IgG-Fc (ms) Affinity Matrix (Fisher) to bind IgG at 4°C for 72 hours on a rotator. Samples with IgG bound to resin were centrifuged at 4,000 rpm and supernatant was collected. IgG samples were washed 3 times with PBS followed by centrifugation to remove supernatant. Samples were buffer exchanged into buffer containing 100mM Tris, 2mM EDTA, and 10mM L-cysteine through centrifugation with Amicon filters, then incubated with papain for 4 hours at 37°C shaking at 80 rpm. The digestion reactions were quenched with 50mM iodoacetamide, buffer exchanged to TBS, and separated by size-exclusion chromatography (SEC) with a Superdex 200 increase 10/300 column (GE Healthcare). Fab and undigested IgG were collected and concentrated and 500 μg Fab was complexed with 10 μg HA for 18 hours at room temperature. Reactions were purified by SEC and immune complexes were collected and concentrated. Negative stain EM grids were prepared as described above.

#### Membrane-bound HA and mAb staining

HEK293T cells were plated into a 6-well plate and transfected overnight with 0.2 μg of plasmid and 10 μg/ml PEI. After 12-16 hours, media was replaced with PFHM-II and cells were rested for 3 days. Transfected cells were trypsinized, washed, and aliquoted. Cells were stained with 10 μg/ml of individual mAbs for 30 minutes. Cells were washed and stained with anti-human IgG Fc-BV421 for 30 minutes. Cells were washed 2 times and run on a BD LSRFortessa X-20. Data were analyzed using FlowJo v10.

#### Single-cell RNA-seq and repertoire analysis

cH5/1^+^ memory B cells (CD19^+^CD27^+^HA^+^) were bulk sorted and partitioned into nanoliter-scale Gel Bead-In-Emulsions (GEMs) to achieve single cell resolution using the 10x Genomics Chromium Controller and according to the manufacturer’s instruction (10x Genomics). The sorted single cells were processed according to 5’ gene expression and B cell Immunoglobulin (Ig) enrichment instruction to prepare the libraries for sequencing. Libraries were sequenced using an Illumina HiSeq 4000 at Northwestern University or an Illumina NextSeq 500 at the University of Chicago. Cellranger Single-Cell Software Suite (version 3.0) was used to perform sample de-multiplexing, barcode processing, and single-cell 5’ and V(D)J counting, and Cellranger mkfastq was used to de-multiplex raw base call (BCL) files into sample-specific fastq files. Subsequently, reads were aligned to the GRCh38 human genome. Cellranger counts and Cellranger vdj package were used to identify gene expression and assemble V(D)J pairs of antibodies.

Single cell datasets were analyzed using Seurat 3 toolkit. We performed conventional pre-process steps for all 22 subjects including cell quality control (QC), normalization, identification of highly variable features, data scaling, and linear dimensional reduction. More specifically, we only kept cells with more than 200 and less then 2500 detected genes for QC step. We normalized the RNA data using conventional log normalization. We identified 2000 highly variable genes for each dataset and performed principle component analysis (PCA) in linear dimensional reduction step. We then integrated all 22 single cell datasets from vaccinated subjects to remove batch effects. In this analysis, we filtered our dataset and only kept cells with both transcriptome and full length and paired heavy and light chain V(D)J sequences (n=1955). From these cells, we identified a group of “VH1-69/Kappa” B cells that used the VH1-69 gene and kappa light chain. We also identified a group of “anchor epitope” B cells by the following rules: 1) VH locus: VH3-23, VH3-30, VH3-30-3, or VH-3-48; 2) VK locus: VK3-11 or VK3-15; 3) JK locus: JK4 or JK5; 4) K-CDR3 length equal to 10; 5) a “NWP” pattern in K-CDR3 peptide.

#### HA conservation modeling

To generate the group 1 HA conservation model, we selected one representative sequence for each group 1 HA subtype from FluDB (https://www.fludb.org/; Table S5) according to a prior study (Burke and Smith, 2014). A multiple sequence alignment from these HA protein sequences was generated using MUSCLE (Edgar, 2004) and the conservation of each residue was quantified using an entropy model (Crooks et al., 2004). HA conservation was visualized on a H1 protein (PDB: 4JTV) using PyMOL (Schrodinger).

#### Statistical analysis

All statistical analyses were performed using Prism software (Graphpad Version 7.0) or *R*. Sample sizes (n) for the number of mAbs tested are indicated in corresponding figures or in the center of pie graphs. Number of biological repeats for experiments and specific tests for statistical significance used are indicated in the corresponding figure legends. *P* values less than or equal to 0.05 were considered significant. * *P* ≤ 0.05, ** *P* ≤ 0.01, *** *P* ≤ 0.001, **** *P* < 0.0001.

## References

Andrews, S.F., Huang, Y., Kaur, K., Popova, L.I., Ho, I.Y., Pauli, N.T., Henry Dunand, C.J., Taylor, W.M., Lim, S., Huang, M., et al. (2015). Immune history profoundly affects broadly protective B cell responses to influenza. Sci Transl Med 7, 316ra192.

Aydillo, T., Escalera, A., Strohmeier, S., Aslam, S., Sanchez-Cespedes, J., Ayllon, J., Roca-Oporto, C., Perez-Romero, P., Montejo, M., Gavalda, J., et al. (2020). Pre-existing Hemagglutinin Stalk Antibodies Correlate with Protection of Lower Respiratory Symptoms in Flu-Infected Transplant Patients. Cell Rep Med 1, 100130.

Bajic, G., Maron, M.J., Adachi, Y., Onodera, T., McCarthy, K.R., McGee, C.E., Sempowski, G.D., Takahashi, Y., Kelsoe, G., Kuraoka, M., et al. (2019). Influenza Antigen Engineering Focuses Immune Responses to a Subdominant but Broadly Protective Viral Epitope. Cell Host Microbe 25, 827–835 e826.

Benton, D.J., Nans, A., Calder, L.J., Turner, J., Neu, U., Lin, Y.P., Ketelaars, E., Kallewaard, N.L., Corti, D., Lanzavecchia, A., et al. (2018). Influenza hemagglutinin membrane anchor. Proc Natl Acad Sci U S A 115, 10112–10117.

Bernstein, D.I., Guptill, J., Naficy, A., Nachbagauer, R., Berlanda-Scorza, F., Feser, J., Wilson, P.C., Solorzano, A., Van der Wielen, M., Walter, E.B., et al. (2020). Immunogenicity of chimeric haemagglutinin-based, universal influenza virus vaccine candidates: interim results of a randomised, placebo-controlled, phase 1 clinical trial. Lancet Infect Dis 20, 80–91.

Bloom, J.D. (2015). Software for the analysis and visualization of deep mutational scanning data. BMC Bioinformatics 16, 168.

Boyoglu-Barnum, S., Hutchinson, G.B., Boyington, J.C., Moin, S.M., Gillespie, R.A., Tsybovsky, Y., Stephens, T., Vaile, J.R., Lederhofer, J., Corbett, K.S., et al. (2020). Glycan repositioning of influenza hemagglutinin stem facilitates the elicitation of protective cross-group antibody responses. Nat Commun 11, 791.

Broecker, F., Liu, S.T.H., Suntronwong, N., Sun, W., Bailey, M.J., Nachbagauer, R., Krammer, F., and Palese, P. (2019). A mosaic hemagglutinin-based influenza virus vaccine candidate protects mice from challenge with divergent H3N2 strains. NPJ Vaccines 4, 31.

Burke, D.F., and Smith, D.J. (2014). A recommended numbering scheme for influenza A HA subtypes. PLoS One 9, e112302.

Cardoso, R.M., Zwick, M.B., Stanfield, R.L., Kunert, R., Binley, J.M., Katinger, H., Burton, D.R., and Wilson, I.A. (2005). Broadly neutralizing anti-HIV antibody 4E10 recognizes a helical conformation of a highly conserved fusion-associated motif in gp41. Immunity 22, 163–173.

Chen, Y.Q., Wohlbold, T.J., Zheng, N.Y., Huang, M., Huang, Y., Neu, K.E., Lee, J., Wan, H., Rojas, K.T., Kirkpatrick, E., et al. (2018). Influenza Infection in Humans Induces Broadly Cross-Reactive and Protective Neuraminidase-Reactive Antibodies. Cell 173, 417–429 e410.

Clark, A.M., DeDiego, M.L., Anderson, C.S., Wang, J., Yang, H., Nogales, A., Martinez-Sobrido, L., Zand, M.S., Sangster, M.Y., and Topham, D.J. (2017). Antigenicity of the 2015-2016 seasonal H1N1 human influenza virus HA and NA proteins. PLoS One 12, e0188267.

Corti, D., Suguitan, A.L., Jr., Pinna, D., Silacci, C., Fernandez-Rodriguez, B.M., Vanzetta, F., Santos, C., Luke, C.J., Torres-Velez, F.J., Temperton, N.J., et al. (2010). Heterosubtypic neutralizing antibodies are produced by individuals immunized with a seasonal influenza vaccine. J Clin Invest 120, 1663–1673.

Cotter, C.R., Jin, H., and Chen, Z. (2014). A single amino acid in the stalk region of the H1N1pdm influenza virus HA protein affects viral fusion, stability and infectivity. PLoS Pathog 10, e1003831.

Crooks, G.E., Hon, G., Chandonia, J.M., and Brenner, S.E. (2004). WebLogo: a sequence logo generator. Genome Res 14, 1188–1190.

Doud, M.B., and Bloom, J.D. (2016). Accurate Measurement of the Effects of All Amino-Acid Mutations on Influenza Hemagglutinin. Viruses 8.

Doud, M.B., Hensley, S.E., and Bloom, J.D. (2017). Complete mapping of viral escape from neutralizing antibodies. PLoS Pathog 13, e1006271.

Dreyfus, C., Laursen, N.S., Kwaks, T., Zuijdgeest, D., Khayat, R., Ekiert, D.C., Lee, J.H., Metlagel, Z., Bujny, M.V., Jongeneelen, M., et al. (2012). Highly conserved protective epitopes on influenza B viruses. Science 337, 1343–1348.

Dugan, H.L., Guthmiller, J.J., Arevalo, P., Huang, M., Chen, Y.Q., Neu, K.E., Henry, C., Zheng, N.Y., Lan, L.Y., Tepora, M.E., et al. (2020). Preexisting immunity shapes distinct antibody landscapes after influenza virus infection and vaccination in humans. Sci Transl Med 12.

Edgar, R.C. (2004). MUSCLE: multiple sequence alignment with high accuracy and high throughput. Nucleic Acids Res 32, 1792–1797.

Eggink, D., Goff, P.H., and Palese, P. (2014). Guiding the immune response against influenza virus hemagglutinin toward the conserved stalk domain by hyperglycosylation of the globular head domain. J Virol 88, 699–704.

Ekiert, D.C., Bhabha, G., Elsliger, M.A., Friesen, R.H., Jongeneelen, M., Throsby, M., Goudsmit, J., and Wilson, I.A. (2009). Antibody recognition of a highly conserved influenza virus epitope. Science 324, 246–251.

Ekiert, D.C., Kashyap, A.K., Steel, J., Rubrum, A., Bhabha, G., Khayat, R., Lee, J.H., Dillon, M.A., O’Neil, R.E., Faynboym, A.M., et al. (2012). Cross-neutralization of influenza A viruses mediated by a single antibody loop. Nature 489, 526–532.

Ellebedy, A.H., Krammer, F., Li, G.M., Miller, M.S., Chiu, C., Wrammert, J., Chang, C.Y., Davis, C.W., McCausland, M., Elbein, R., et al. (2014). Induction of broadly cross-reactive antibody responses to the influenza HA stem region following H5N1 vaccination in humans. Proc Natl Acad Sci U S A 111, 13133–13138.

Gallagher, J.R., McCraw, D.M., Torian, U., Gulati, N.M., Myers, M.L., Conlon, M.T., and Harris, A.K. (2018). Characterization of Hemagglutinin Antigens on Influenza Virus and within Vaccines Using Electron Microscopy. Vaccines (Basel) 6.

Guthmiller, J.J., Dugan, H.L., Neu, K.E., Lan, L.Y., and Wilson, P.C. (2019). An Efficient Method to Generate Monoclonal Antibodies from Human B Cells. Methods Mol Biol 1904, 109–145.

Guthmiller, J.J., Lan, L.Y., Fernandez-Quintero, M.L., Han, J., Utset, H.A., Bitar, D.J., Hamel, N.J., Stovicek, O., Li, L., Tepora, M., et al. (2020). Polyreactive Broadly Neutralizing B cells Are Selected to Provide Defense against Pandemic Threat Influenza Viruses. Immunity.

Hai, R., Krammer, F., Tan, G.S., Pica, N., Eggink, D., Maamary, J., Margine, I., Albrecht, R.A., and Palese, P. (2012). Influenza viruses expressing chimeric hemagglutinins: globular head and stalk domains derived from different subtypes. J Virol 86, 5774–5781.

Han, J., Schmitz, A.J., Richey, S.T., Dai, Y.-N., Turner, H.L., Mohammed, B.M., Fremont, D.H., Ellebedy, A.H., and Ward, A.B. (2020). Polyclonal epitope cartography reveals the temporal dynamics and diversity of human antibody responses to H5N1 vaccination. bioRxiv, 2020.2006.2016.155754.

Henry, C., Zheng, N.Y., Huang, M., Cabanov, A., Rojas, K.T., Kaur, K., Andrews, S.F., Palm, A.E., Chen, Y.Q., Li, Y., et al. (2019). Influenza Virus Vaccination Elicits Poorly Adapted B Cell Responses in Elderly Individuals. Cell Host Microbe 25, 357–366 e356.

Henry Dunand, C.J., Leon, P.E., Huang, M., Choi, A., Chromikova, V., Ho, I.Y., Tan, G.S., Cruz, J., Hirsh, A., Zheng, N.Y., et al. (2016). Both Neutralizing and Non-Neutralizing Human H7N9 Influenza Vaccine-Induced Monoclonal Antibodies Confer Protection. Cell Host Microbe 19, 800–813.

Henry Dunand, C.J., Leon, P.E., Kaur, K., Tan, G.S., Zheng, N.Y., Andrews, S., Huang, M., Qu, X., Huang, Y., Salgado-Ferrer, M., et al. (2015). Preexisting human antibodies neutralize recently emerged H7N9 influenza strains. J Clin Invest 125, 1255–1268.

Impagliazzo, A., Milder, F., Kuipers, H., Wagner, M.V., Zhu, X., Hoffman, R.M., van Meersbergen, R., Huizingh, J., Wanningen, P., Verspuij, J., et al. (2015). A stable trimeric influenza hemagglutinin stem as a broadly protective immunogen. Science 349, 1301–1306.

Joyce, M.G., Wheatley, A.K., Thomas, P.V., Chuang, G.Y., Soto, C., Bailer, R.T., Druz, A., Georgiev, I.S., Gillespie, R.A., Kanekiyo, M., et al. (2016). Vaccine-Induced Antibodies that Neutralize Group 1 and Group 2 Influenza A Viruses. Cell 166, 609–623.

Kirkpatrick, E., Qiu, X., Wilson, P.C., Bahl, J., and Krammer, F. (2018). The influenza virus hemagglutinin head evolves faster than the stalk domain. Sci Rep 8, 10432.

Krammer, F., Pica, N., Hai, R., Margine, I., and Palese, P. (2013). Chimeric hemagglutinin influenza virus vaccine constructs elicit broadly protective stalk-specific antibodies. J Virol 87, 6542–6550.

Krystal, M., Elliott, R.M., Benz, E.W., Jr., Young, J.F., and Palese, P. (1982). Evolution of influenza A and B viruses: conservation of structural features in the hemagglutinin genes. Proc Natl Acad Sci U S A 79, 4800–4804.

Lander, G.C., Stagg, S.M., Voss, N.R., Cheng, A., Fellmann, D., Pulokas, J., Yoshioka, C., Irving, C., Mulder, A., Lau, P.W., et al. (2009). Appion: an integrated, database-driven pipeline to facilitate EM image processing. J Struct Biol 166, 95–102.

Li, G.M., Chiu, C., Wrammert, J., McCausland, M., Andrews, S.F., Zheng, N.Y., Lee, J.H., Huang, M., Qu, X., Edupuganti, S., et al. (2012). Pandemic H1N1 influenza vaccine induces a recall response in humans that favors broadly cross-reactive memory B cells. Proc Natl Acad Sci U S A 109, 9047–9052.

Linderman, S.L., Chambers, B.S., Zost, S.J., Parkhouse, K., Li, Y., Herrmann, C., Ellebedy, A.H., Carter, D.M., Andrews, S.F., Zheng, N.Y., et al. (2014). Potential antigenic explanation for atypical H1N1 infections among middle-aged adults during the 2013-2014 influenza season. Proc Natl Acad Sci U S A 111, 15798–15803.

Liu, S.T.H., Behzadi, M.A., Sun, W., Freyn, A.W., Liu, W.C., Broecker, F., Albrecht, R.A., Bouvier, N.M., Simon, V., Nachbagauer, R., et al. (2018). Antigenic sites in influenza H1 hemagglutinin display species-specific immunodominance. J Clin Invest 128, 4992–4996.

Mena, I., Nelson, M.I., Quezada-Monroy, F., Dutta, J., Cortes-Fernandez, R., Lara-Puente, J.H., Castro-Peralta, F., Cunha, L.F., Trovao, N.S., Lozano-Dubernard, B., et al. (2016). Origins of the 2009 H1N1 influenza pandemic in swine in Mexico. Elife 5.

Nachbagauer, R., Feser, J., Naficy, A., Bernstein, D.I., Guptill, J., Walter, E.B., Berlanda-Scorza, F., Stadlbauer, D., Wilson, P.C., Aydillo, T., et al. (2020). A chimeric hemagglutinin-based universal influenza virus vaccine approach induces broad and long-lasting immunity in a randomized, placebo-controlled phase I trial. Nat Med.

Nachbagauer, R., Wohlbold, T.J., Hirsh, A., Hai, R., Sjursen, H., Palese, P., Cox, R.J., and Krammer, F. (2014). Induction of broadly reactive anti-hemagglutinin stalk antibodies by an H5N1 vaccine in humans. J Virol 88, 13260–13268.

Ng, S., Nachbagauer, R., Balmaseda, A., Stadlbauer, D., Ojeda, S., Patel, M., Rajabhathor, A., Lopez, R., Guglia, A.F., Sanchez, N., et al. (2019). Novel correlates of protection against pandemic H1N1 influenza A virus infection. Nat Med 25, 962–967.

Ofek, G., Tang, M., Sambor, A., Katinger, H., Mascola, J.R., Wyatt, R., and Kwong, P.D. (2004). Structure and mechanistic analysis of the anti-human immunodeficiency virus type 1 antibody 2F5 in complex with its gp41 epitope. J Virol 78, 10724–10737.

Ohmit, S.E., Petrie, J.G., Cross, R.T., Johnson, E., and Monto, A.S. (2011). Influenza hemagglutination-inhibition antibody titer as a correlate of vaccine-induced protection. J Infect Dis 204, 1879–1885.

Park, J.K., Xiao, Y., Ramuta, M.D., Rosas, L.A., Fong, S., Matthews, A.M., Freeman, A.D., Gouzoulis, M.A., Batchenkova, N.A., Yang, X., et al. (2020). Pre-existing immunity to influenza virus hemagglutinin stalk might drive selection for antibody-escape mutant viruses in a human challenge model. Nat Med 26, 1240–1246.

Pettersen, E.F., Goddard, T.D., Huang, C.C., Couch, G.S., Greenblatt, D.M., Meng, E.C., and Ferrin, T.E. (2004). UCSF Chimera--a visualization system for exploratory research and analysis. J Comput Chem 25, 1605–1612.

Pica, N., and Palese, P. (2013). Toward a universal influenza virus vaccine: prospects and challenges. Annu Rev Med 64, 189–202.

Raymond, D.D., Bajic, G., Ferdman, J., Suphaphiphat, P., Settembre, E.C., Moody, M.A., Schmidt, A.G., and Harrison, S.C. (2018). Conserved epitope on influenza-virus hemagglutinin head defined by a vaccine-induced antibody. Proc Natl Acad Sci U S A 115, 168–173.

Scheres, S.H. (2012). RELION: implementation of a Bayesian approach to cryo-EM structure determination. J Struct Biol 180, 519–530.

Smith, K., Garman, L., Wrammert, J., Zheng, N.Y., Capra, J.D., Ahmed, R., and Wilson, P.C. (2009). Rapid generation of fully human monoclonal antibodies specific to a vaccinating antigen. Nat Protoc 4, 372–384.

Sui, J., Hwang, W.C., Perez, S., Wei, G., Aird, D., Chen, L.M., Santelli, E., Stec, B., Cadwell, G., Ali, M., et al. (2009). Structural and functional bases for broad-spectrum neutralization of avian and human influenza A viruses. Nat Struct Mol Biol 16, 265–273.

Suloway, C., Pulokas, J., Fellmann, D., Cheng, A., Guerra, F., Quispe, J., Stagg, S., Potter, C.S., and Carragher, B. (2005). Automated molecular microscopy: the new Leginon system. J Struct Biol 151, 41–60.

Sun, W., Kirkpatrick, E., Ermler, M., Nachbagauer, R., Broecker, F., Krammer, F., and Palese, P. (2019). Development of Influenza B Universal Vaccine Candidates Using the “Mosaic” Hemagglutinin Approach. J Virol 93.

van der Lubbe, J.E.M., Huizingh, J., Verspuij, J.W.A., Tettero, L., Schmit-Tillemans, S.P.R., Mooij, P., Mortier, D., Koopman, G., Bogers, W., Dekking, L., et al. (2018). Mini-hemagglutinin vaccination induces cross-reactive antibodies in pre-exposed NHP that protect mice against lethal influenza challenge. NPJ Vaccines 3, 25.

Wasilewski, S., Calder, L.J., Grant, T., and Rosenthal, P.B. (2012). Distribution of surface glycoproteins on influenza A virus determined by electron cryotomography. Vaccine 30, 7368–7373.

Weidenbacher, P.A., and Kim, P.S. (2019). Protect, modify, deprotect (PMD): A strategy for creating vaccines to elicit antibodies targeting a specific epitope. Proc Natl Acad Sci U S A 116, 9947–9952.

Whittle, J.R., Zhang, R., Khurana, S., King, L.R., Manischewitz, J., Golding, H., Dormitzer, P.R., Haynes, B.F., Walter, E.B., Moody, M.A., et al. (2011). Broadly neutralizing human antibody that recognizes the receptor-binding pocket of influenza virus hemagglutinin. Proc Natl Acad Sci U S A 108, 14216–14221.

Wrammert, J., Koutsonanos, D., Li, G.M., Edupuganti, S., Sui, J., Morrissey, M., McCausland, M., Skountzou, I., Hornig, M., Lipkin, W.I., et al. (2011). Broadly cross-reactive antibodies dominate the human B cell response against 2009 pandemic H1N1 influenza virus infection. J Exp Med 208, 181–193.

Wrammert, J., Smith, K., Miller, J., Langley, W.A., Kokko, K., Larsen, C., Zheng, N.Y., Mays, I., Garman, L., Helms, C., et al. (2008). Rapid cloning of high-affinity human monoclonal antibodies against influenza virus. Nature 453, 667–671.

Wu, N.C., Thompson, A.J., Lee, J.M., Su, W., Arlian, B.M., Xie, J., Lerner, R.A., Yen, H.L., Bloom, J.D., and Wilson, I.A. (2020). Different genetic barriers for resistance to HA stem antibodies in influenza H3 and H1 viruses. Science 368, 1335–1340.

Wu, N.C., and Wilson, I.A. (2020). Influenza Hemagglutinin Structures and Antibody Recognition. Cold Spring Harb Perspect Med 10.

Yassine, H.M., Boyington, J.C., McTamney, P.M., Wei, C.J., Kanekiyo, M., Kong, W.P., Gallagher, J.R., Wang, L., Zhang, Y., Joyce, M.G., et al. (2015). Hemagglutinin-stem nanoparticles generate heterosubtypic influenza protection. Nat Med 21, 1065–1070.

