## Supplement Figures, Tables, and References for "A public broadly neutralizing antibody class targets a membrane-proximal anchor epitope of influenza virus hemagglutinin"

Figure S1

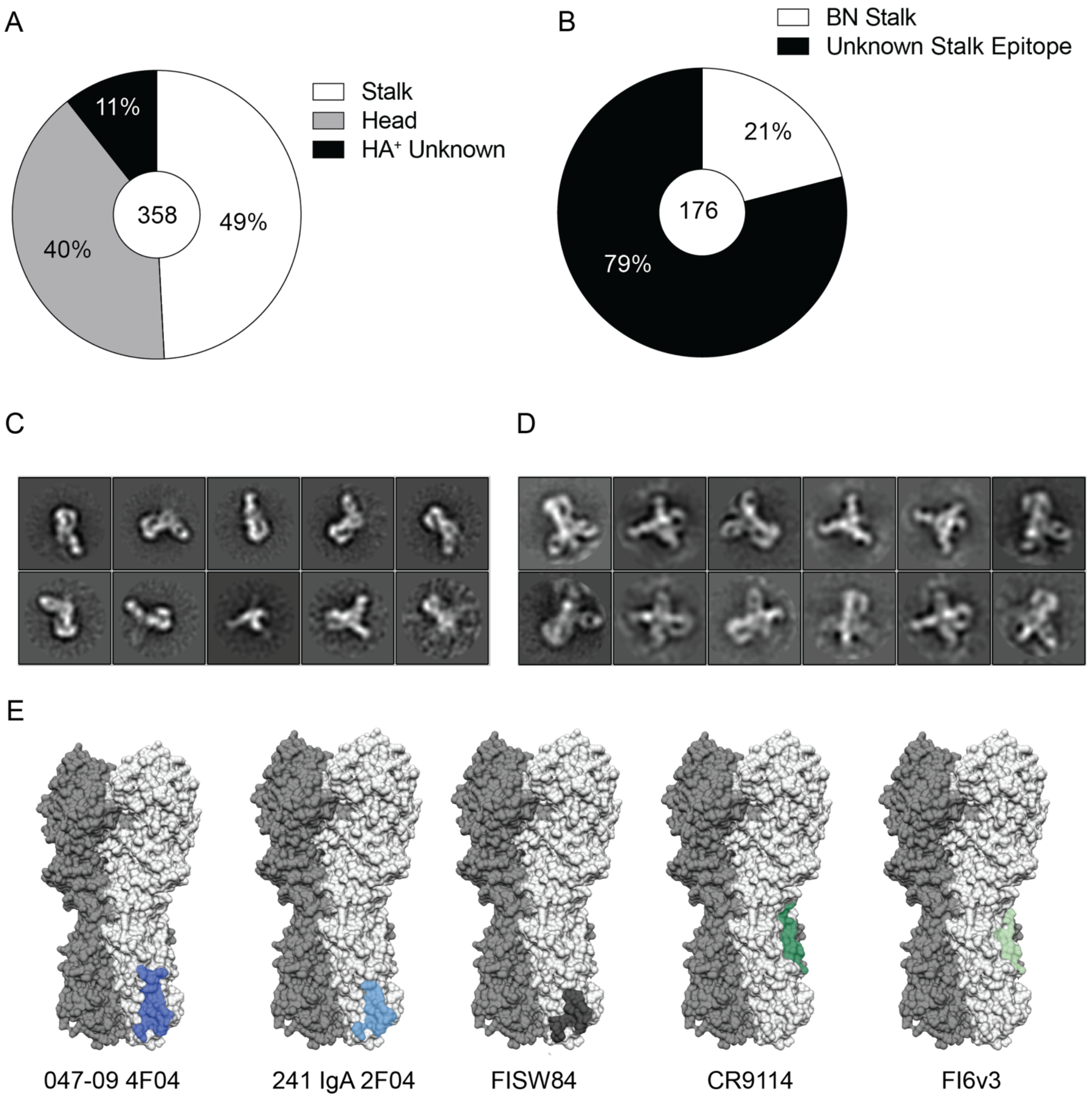


**Figure S1: Binding features of isolated mAbs and 2D class average of anchor epitope binding mAbs. Related to Figure 1.** (**A**) Proportion of HA$^+$ mAbs binding to distinct HA domains. (**B**) Proportion of stalk-binding mAbs binding the BN stalk domain. (**C**-**D**) Negative stain 2D class averages of 047-09 4F04 (**C**) and 241 IgA 2F04 (**D**) binding A/California/4/2009 HA. (**E**) Binding footprint of mAbs binding the anchor epitope (047-09 4F04, 241 IgA 2F04, and FISW84) and the BN stalk epitope (CR9114 and FI6v3) on A/California/7/2009.

Figure S2

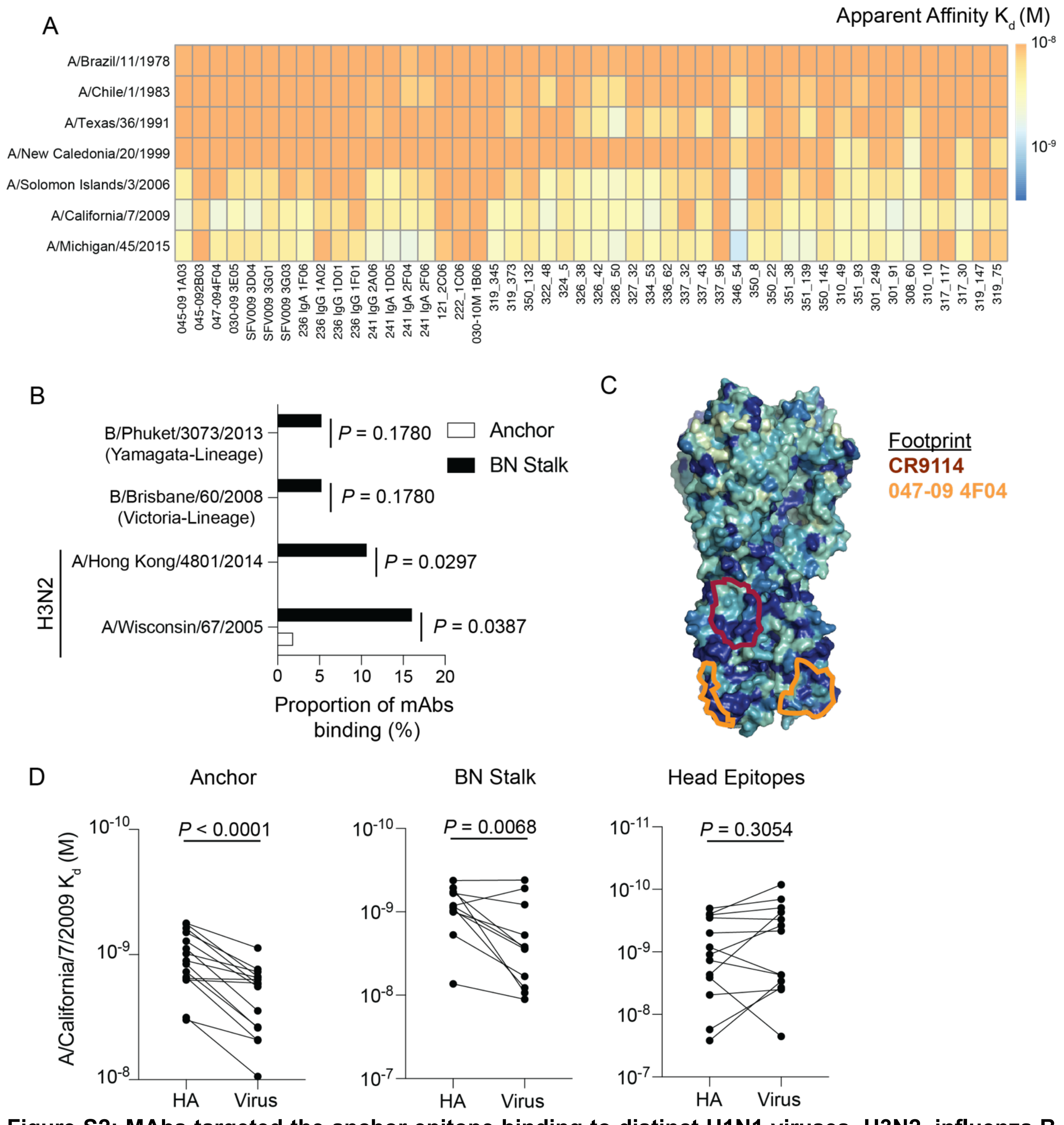


**Figure S2: MAbs targeted the anchor epitope binding to distinct H1N1 viruses, H3N2, influenza B viruses and comparison of mAbs binding virus versus HA. Related to Figure 2.** (**A**) Heatmap of apparent affinity ($K_d$; M) of anchor mAb binding to historical and recent H1N1 viruses. (**B**) Proportion of anchor binding and BN stalk binding mAbs binding to influenza B viruses and H3N2 viruses. (**C**) Conservation of group 1 influenza virus HAs, as shown on A/California/04/2009 (PDB: 4JTV), with the binding footprint of CR9114 (maroon) and 047-09 4F04 (orange) outlined. (**D**) Apparent affinity ($K_d$; M) of mAbs binding anchor, BN stalk, or head epitope binding to recombinant HA from A/California/7/2009 H1N1 and A/California/7/2009 virus. Lines connect the same mAb. Data in **B** were analyzed by Fisher's Exact tests and data in **D** were analyzed using a paired non-parametric Wilcoxon matched-pairs signed rank test.

Figure S3

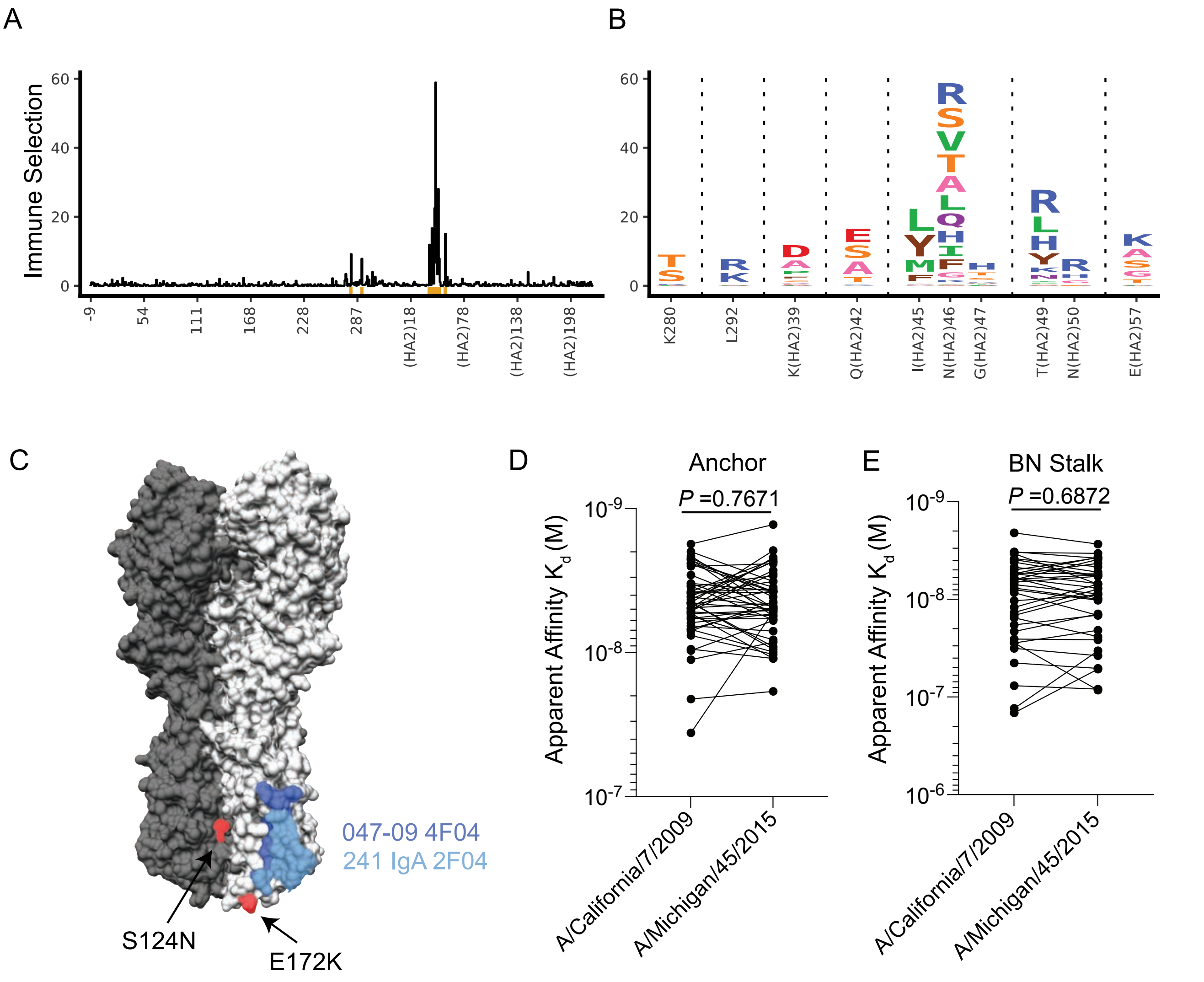


**Figure S3: Deep mutational scanning of 045-09 2B06 and mAb affinity for A/Michigan/45/2015 H1N1. Related to Figure 2.** (**A**-**B**) 045-09 2B06 was subjected to deep mutational scanning with a mutant library of A/WSN/1933 (H1N1). (**A**) Differential selection at each site in HA (H3 numbering). (**B**) Logo plot of key sites of escape, with the height of each letter proportional to the antibody selection for mutations to that amino acid at that site. (**C**) Differences in mutations found on the stalk domain of A/Michigan/45/2015 relative to the binding footprints of 047-09 4F04 and 241 IgA 2F04. (**D**-**E**) Apparent Affinity ($K_d$; M) of anchor binding mAbs (**D**) and BN stalk binding mAbs (**E**) to A/California/7/2009 and A/Michigan/45/2009. Lines connect the same mAb. Data in **D** and **E** were analyzed using a paired non-parametric Wilcoxon matched-pairs signed rank test.

Figure S4

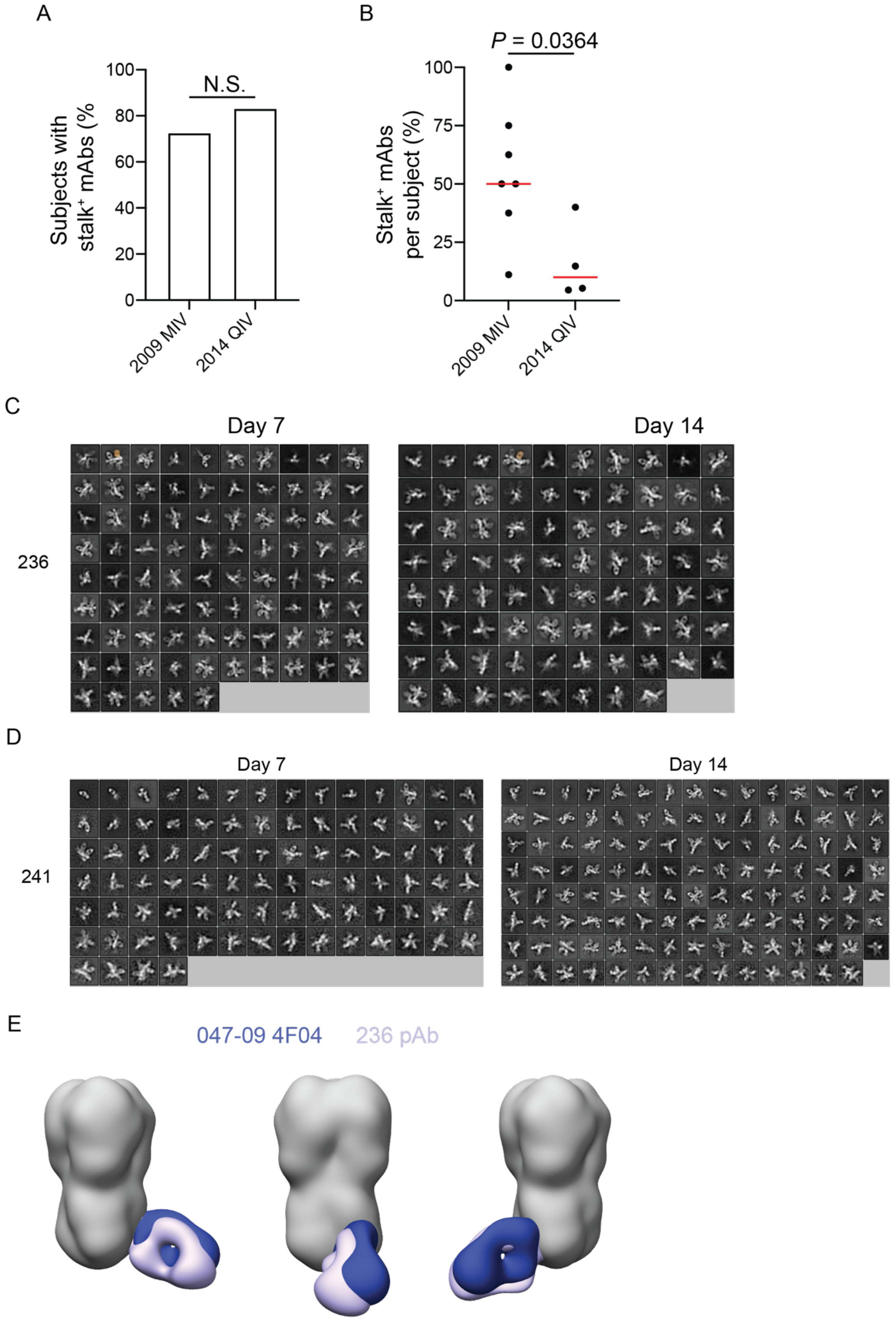

A
Subjects with stalk⁺ mAbs (%
100
80
60
40
20
0
N.S.
2009 MIV
2014 QIV
B
P = 0.0364
Stalk⁺ mAbs per subject (%)
100
75
50
25
0
2009 MIV
2014 QIV
C
Day 7
Day 14
236
D
Day 7
Day 14
241
E
047-09 4F04
236 pAb

**Figure S4: Characterization of stalk binding mAbs induced by influenza virus vaccines. Related to Figure 4.** (**A**) Proportion of subjects with stalk$^+$ mAbs from the 2009 MIV or 2014 QIV cohorts. (**B**) Proportion of mAbs targeting the stalk per person in the 2009 MIV and 2014 QIV cohorts. Red line represents the median. Each symbol in **B** represents one subject. Only subjects with stalk$^+$ mAbs were included in the analysis. (**C**-**D**) 2D class averages of pAbs from subject 236 (**C**) and 241 (**D**) at days 7 and 14 post immunization binding to A/Michigan/45/2015 HA. (**E**) 3D reconstruction of the overlapping of 047-09 4F04, 236 pAb, and FISW84. Data in **A** were analyzed by Fisher's Exact tests and data in **B** were analyzed by unpaired non-parametric Mann-Whitney test.

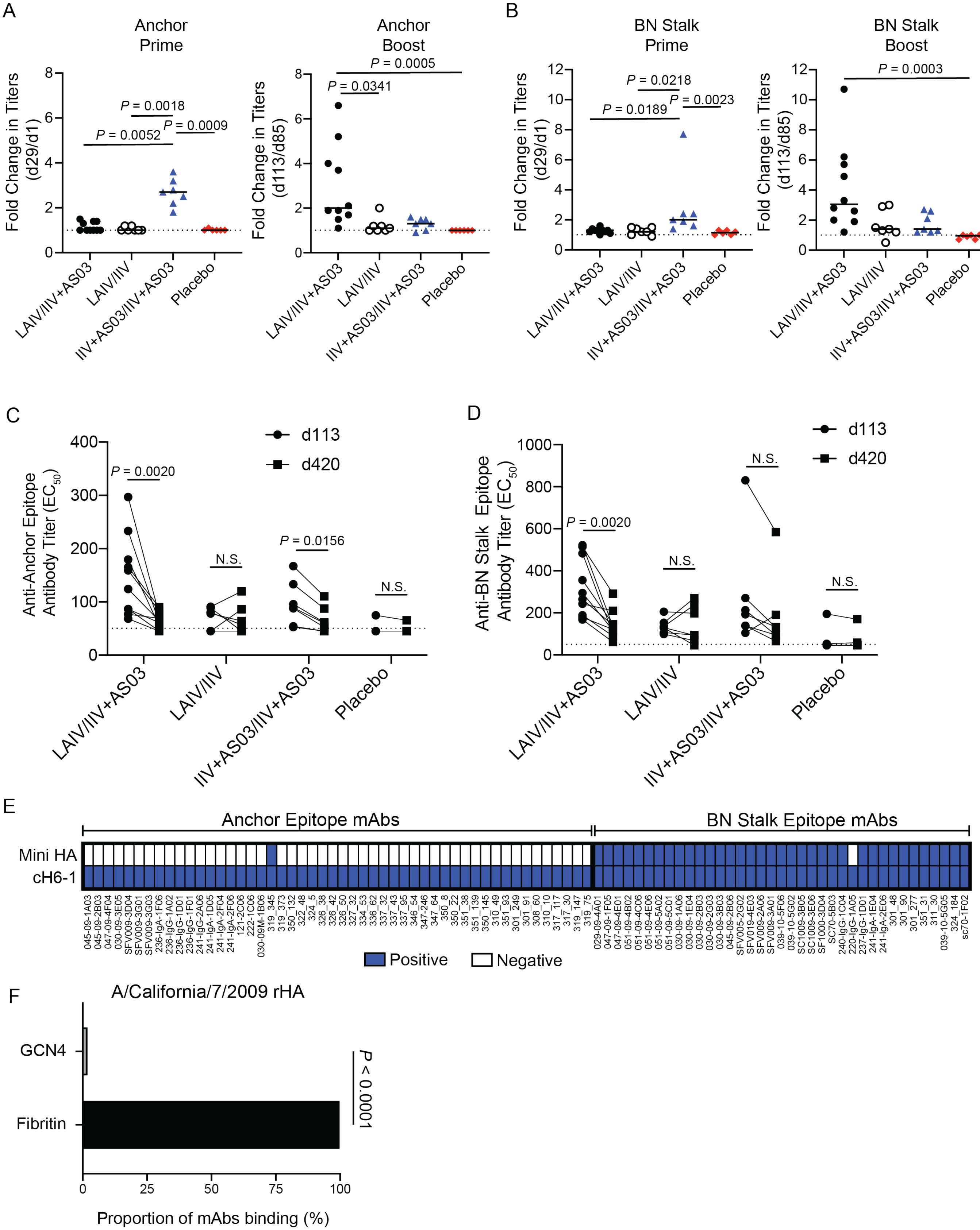


**Figure S5: Seroconversion against anchor and BN stalk epitopes after cHA vaccination and mAb binding to various recombinant HAs. Related to Figure 5.** (**A-B**) Seroconversion (d28/d0) of antibodies competing for binding against the anchor epitope (**A**) and the BN stalk epitope (**B**). Bar represents the median and each symbol represents one subject. (**C**-**D**) Antibody titers ($EC_{50}$) of serum antibodies collected on day 112 and day 420 against the anchor epitope (**C**) and the BN stalk epitope (**D**). Lines connect titers from the same subject and each pair of symbols represents one subject. (**E**) Anchor and BN stalk targeting mAbs binding to mini-HA and cH6/1. (**F**) Proportion of anchor epitope targeting mAbs binding to A/California/7/2009 recombinant HA with a GCN4 or fibritin trimerization domain. Data in **A** and **B** were analyzed by non-parametric Kruskal-Wallis Tests, data in **C** and **D** were analyzed using a paired non-parametric Wilcoxon matched-pairs signed rank test, and data in **F** were analyzed by a Fisher's Exact test.

Figure S6

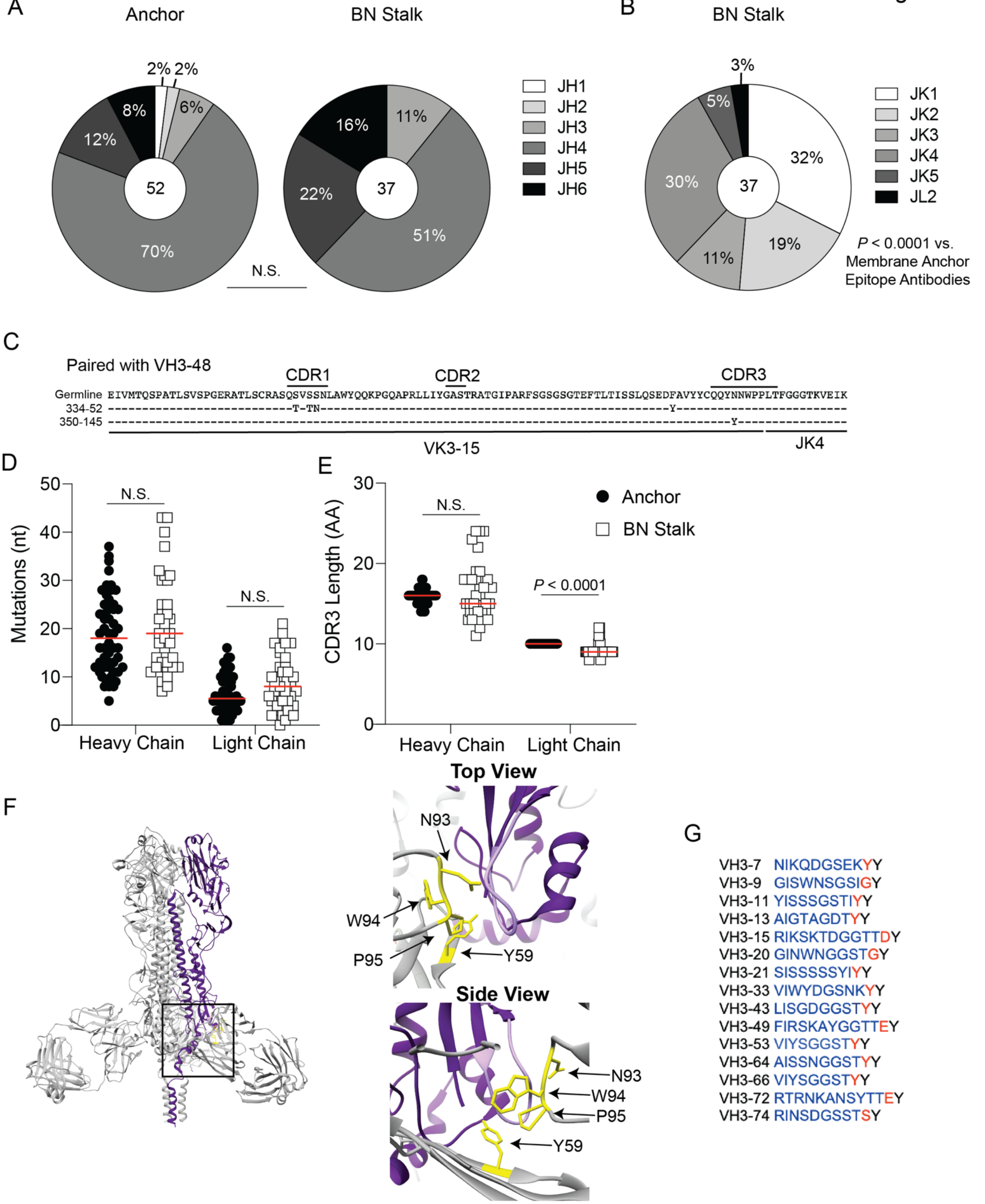


**Figure S6: Additional repertoire features of mAs binding the anchor epitope. Related to Figure 6.** (**A**) JH gene usage by anchor and BN stalk binding antibodies. (**B**) JK gene usage of mAbs binding the BN stalk epitope. (**C**) VJ sequences of kappa chains of the public clone paired VH3-48. (**D-E**) Mutations (**D**) and CDR3 amino acid (AA) lengths (**E**) of heavy and light chains of mAbs binding the anchor or BN stalk epitopes. Red bar represents median. (**F**) NWP motif of K-CDR3 and Y59 following H-CDR2 (**H**) of FISW84 (PDB: 6HJQ). (**G**) H-CDR2 sequences of germline VH3 genes. H-CDR2 is highlighted in blue and the first amino acid following H-CDR2 is highlighted in red. Data in **A** were analyzed using a Chi-square test, and data in **D** and **E** were analyzed by unpaired non-parametric Mann-Whitney tests.

**Table S1: Subject information and demographics.**

| Subject | Vaccine/Infection | Age | Sex |
|---|---|---|---|
| 029-09 | 2009 MIV | 24 | M |
| 030-09 | 2009 MIV | 31 | F |
| 045-09 | 2009 MIV | 24 | M |
| 047-09 | 2009 MIV | 22 | M |
| 051-09 | 2009 MIV | 42 | M |
| SFV005 | 2009 MIV | 60 | F |
| SFV009 | 2009 MIV | 31 | F |
| SFV015 | 2009 MIV | 48 | F |
| SFV018 | 2009 MIV | 58 | F |
| SFV019 | 2009 MIV | 48 | F |
| SFV020 | 2009 MIV | 64 | F |
| sc70 | 2009 pH1N1 Infection | 38 | F |
| sc1009 | 2009 pH1N1 Infection | 21 | M |
| SF1000 | 2009 pH1N1 Infection | 37 | M |
| 008-10 | 2010 TIV | 26 | F |
| 011-10 | 2010 TIV | 30 | M |
| 014-10 | 2010 TIV | 27 | M |
| 017-10 | 2010 TIV | 24 | M |
| 019-10 | 2010 TIV | 22 | F |
| 028-10 | 2010 TIV | 38 | M |
| 034-10 | 2010 TIV | 39 | F |
| 039-10 | 2010 TIV | 25 | M |
| 051-10 | 2010 TIV | 43 | M |
| 220-14 | 2014 QIV | 24 | F |
| 221-14 | 2014 QIV | 34 | F |
| 236-14 | 2014 QIV | 32 | F |
| 237-14 | 2014 QIV | 32 | F |
| 240-14 | 2014 QIV | 28 | M |
| 241-14 | 2014 QIV | 29 | F |
| 121/324 (d7/d112) | cHA – IIV+AS03/IIV+AS03 | 24 | F |
| 222/310 (d91/d112) | cHA – LAIV/IIV+AS03 | 25 | F |
| 301 | cHA – IIV+AS03/IIV+AS03 | 34 | F |
| 308 | cHA – LAIV/IIV+AS03 | 29 | F |
| 311 | cHA – IIV+AS03/IIV+AS03 | 28 | M |
| 317 | cHA – LAIV/IIV+AS03 | 37 | M |
| 319 | cHA – LAIV/IIV+AS03 | 31 | F |
| 322 | cHA – IIV+AS03/IIV+AS03 | 20 | F |
| 326 | cHA – LAIV/IIV | 27 | M |
| 327 | cHA – LAIV/IIV+AS03 | 22 | F |
| 334 | cHA – LAIV/IIV+AS03 | 33 | M |

| | | | |
|---|---|---|---|
| 336 | cHA – LAIV/IIV+AS03 | 36 | F |
| 337 | cHA – LAIV/IIV+AS03 | 27 | F |
| 346 | cHA – LAIV/IIV | 20 | F |
| 347 | cHA – LAIV/IIV+AS03 | 31 | F |
| 350 | cHA – LAIV/IIV+AS03 | 35 | F |
| 351 | cHA – IIV+AS03/IIV+AS03 | 27 | F |

**Table S2: Anchor epitope binding mAb information.**

| mAb | Cellular Source | VH | DH | JH | HC Clone # | VK | JK | KC Clone # |
|---|---|---|---|---|---|---|---|---|
| 030-09 3E05 | Plasmablast | 3-23 | 3-22 | 4 | Non-clonal | 3-15 | 5 | 9 |
| 030-09M 1B06 | MBC | 3-23 | 3-10 | 4 | Non-clonal | 3-11 | 5 | 7 |
| 045-09 1A03 | Plasmablast | 3-23 | 6-13 | 4 | Non-clonal | 3-11 | 4 | 6 |
| 045-09 2B03 | Plasmablast | 3-23 | 5-12 | 4 | Non-clonal | 3-11 | 5 | 7 |
| 047-09 4F04 | Plasmablast | 3-48 | 6-19 | 4 | Non-clonal | 3-15 | 4 | 8 |
| SFV009 3D04 | Plasmablast | 3-23 | 2-2 | 4 | Non-clonal | 3-15 | 4 | 8 |
| SFV009 3G01 | Plasmablast | 3-30 | 3-16 | 4 | Non-clonal | 3-11 | 5 | 7 |
| SFV009 3G03 | Plasmablast | 3-23 | 3-15-3 | 5 | Non-clonal | 3-11 | 5 | 7 |
| 236 IgG 1A02 | Plasmablast | 3-23 | 2-2 | 4 | 2 | 3-11 | 5 | 7 |
| 236 IgG 1D01 | Plasmablast | 3-23 | 6-19 | 4 | Non-clonal | 3-11 | 5 | 7 |
| 236 IgG 1F01 | Plasmablast | 3-23 | 2-2 | 4 | 2 | 3-11 | 5 | 7 |
| 236 IgA 1F06 | Plasmablast | 3-23 | 2-2 | 4 | 2 | 3-11 | 5 | 7 |
| 241 IgG 2A06 | Plasmablast | 3-23 | 6-19 | 5 | Non-Clonal | 3-15 | 4 | 8 |
| 241 IgA 1D05 | Plasmablast | 3-23 | 5-18 | 4 | 3 | 3-15 | 5 | 9 |
| 241 IgA 2F04 | Plasmablast | 3-23 | 2-21 | 4 | 3 | 3-15 | 5 | 9 |
| 241 IgA 2F06 | Plasmablast | 3-23 | 3-9 | 4 | 3 | 3-15 | 5 | 9 |
| 121 2C06 | Plasmablast | 3-23 | 2-15 | 4 | Non-clonal | 3-15 | 4 | 8 |
| 222 1C06 | Plasmablast | 3-48 | 2-15-2 | 4 | Non-clonal | 3-15 | 4 | 8 |
| 301_91 | MBC | 3-23 | 3-9 | 5 | Non-clonal | 3-11 | 5 | 7 |
| 301_249 | MBC | 3-48 | 5-24 | 3 | Non-clonal | 3-11 | 4 | Non-clonal |
| 301_275 | MBC | 3-23 | 2-21 | 4 | 1 | 3-15 | 4 | 8 |
| 308_60 | MBC | 3-23 | 1-26 | 4 | Non-clonal | 3-11 | 4 | 8 |
| 310_10 | MBC | 3-23 | 1-14 | 4 | Non-clonal | 3-15 | 4 | 8 |
| 310_49 | MBC | 3-23 | 3-22 | 4 | Non-clonal | 3-15 | 5 | 9 |
| 317_30 | MBC | 3-23 | 6-13 | 4 | Non-clonal | 3-15 | 4 | 8 |
| 317_117 | MBC | 3-23 | 2-2 | 4 | Non-clonal | 3-15 | 5 | 9 |
| 319_75 | MBC | 3-23 | 3-22 | 3 | Non-clonal | 3-11 | 5 | 7 |
| 319_147 | MBC | 3-48 | 3-15-3 | 6 | Non-clonal | 3-15 | 4 | 8 |
| 319_345 | MBC | 3-23 | 3-9 | 4 | 4 | 3-15 | 4 | 8 |
| 319_373 | MBC | 3-23 | 6-19 | 4 | 4 | 3-15 | 4 | 8 |
| 322_48 | MBC | 3-23 | 1-26 | 4 | 1 | 3-15 | 5 | 9 |
| 324_5 | MBC | 3-23 | 2-21 | 3 | Non-clonal | 3-15 | 5 | 9 |
| 326_38 | MBC | 3-23 | 1-26 | 4 | Non-clonal | 3-11 | 4 | 6 |
| 326_42 | MBC | 3-23 | 3-16 | 5 | Non-clonal | 3-15 | 4 | 8 |
| 326_50 | MBC | 3-30 | 5-12 | 4 | Non-clonal | 3-15 | 5 | 9 |
| 327_32 | MBC | 3-23 | 3-10 | 4 | Non-clonal | 3-15 | 4 | 8 |
| 334_52 | MBC | 3-48 | 6-13 | 6 | 5 | 3-15 | 4 | 8 |
| 334_53 | MBC | 3-23 | 2-2 | 4 | Non-clonal | 3-15 | 5 | 9 |
| 334_62 | MBC | 3-23 | 2-2 | 1 | Non-clonal | 3-11 | 4 | 6 |
| 337_32 | MBC | 3-30 | 5-24 | 4 | Non-clonal | 3-11 | 5 | 7 |
| 337_43 | MBC | 3-23 | 5-24 | 4 | Non-clonal | 3-15 | 4 | 8 |
| 337_95 | MBC | 3-48 | 3-16 | 6 | Non-clonal | 3-15 | 5 | 9 |
| 346_54 | MBC | 3-30-3 | 3-16 | 5 | Non-clonal | 3-11 | 4 | 6 |
| 347_64 | MBC | 3-23 | 1-7 | 2 | Non-clonal | 3-11 | 5 | 7 |
| 347_246 | MBC | 3-23 | 1-1 | 5 | Non-clonal | 3-15 | 5 | 9 |
| 350_8 | MBC | 3-23 | 3-16 | 4 | Non-clonal | 3-15 | 4 | 8 |
| 350_22 | MBC | 3-23 | 6-19 | 4 | Non-clonal | 3-11 | 4 | 6 |
| 350_132 | MBC | 3-48 | 1-26 | 4 | Non-clonal | 3-15 | 4 | 8 |
| 350_145 | MBC | 3-48 | 4-17 | 6 | 5 | 3-15 | 4 | 8 |
| 351_38 | MBC | 3-23 | 6-13 | 4 | Non-clonal | 3-15 | 4 | 8 |
| 351_93 | MBC | 3-23 | 1-1 | 4 | Non-clonal | 3-15 | 4 | 8 |
| 351_139 | MBC | 3-23 | 5-24 | 4 | Non-clonal | 3-11 | 4 | 6 |

**Table S3: Mutation information for Figure 2C-D.**

| Mutation | Naturally Occurring? | 045-09 2B06 Deep Mutational Scan? | Other Experimentally Determined? | Note | Source |
|---|---|---|---|---|---|
| **H38S (HA1)** | | No | Yes | Affects C179 binding | (Doud et al., 2018) |
| **L292R (HA1)** | | Yes | | In footprint of many VH1-69 utilizing mAbs | (Lang et al., 2017) |
| **I323V (HA1)** | Yes | No | | Present in pH1N1 viruses | (Xu et al., 2012) |
| **K39D (HA2)** | | Yes | Yes | Contacts with VH3-30 mAb | (Fu et al., 2016) |
| **Q42E (HA2)** | | Yes | Yes | Interact with CR9114 and FI6v3 | (Wu et al., 2020) |
| **A44V (HA2)** | | No | Yes | Expands in presence of stalk binding mAbs, escape mutant of 6F12 | (Park et al., 2020; Tan et al., 2012) |
| **I45L (HA2)** | | Yes | Yes | Mutation associated with resistance to group 1 neutralizing stalk mAbs | (Lingwood et al., 2012; Wu et al., 2020; Yassine et al., 2018) |
| **D46R (HA2)** | | Yes | | Pre-pH1N1 viruses express N46 | |
| **E47K (HA2)** | Yes | Yes | | Associated with HA protein stability | (Cotter et al., 2014) |
| **T49R (HA2)** | | Yes | | Mutation associated with resistance to group 1 neutralizing stalk mAbs | (Lingwood et al., 2012; Yassine et al., 2018) |
| **N50R (HA2)** | | Yes | | Interacts with CR9114 | (Dreyfus et al., 2012) |
| **V52A (HA2)** | | No | Yes | Affects FI6v3 mAb binding | (Wu et al., 2020) |
| **I56A (HA2)** | | No | Yes | Affects F10 mAb binding | (Sui et al., 2009) |
| **E57K (HA2)** | | Yes | | Critical residue for membrane fusion | (Vanderlinden et al., 2010) |

**Table S4: MAbs used in Figure 3 infection studies.**

| mAb name | Epitope Specificity |
|---|---|
| 241 IgG 2A06 | Anchor |
| 047-09 4F04 | Anchor |
| 030-09 3E05 | Anchor |
| SFV009 3G01 | Anchor |
| 236 IgG 1A02 | Anchor |
| 045-09 2B06 | BN Stalk |
| SFV005 2G02 | BN Stalk |
| SFV019 4E03 | BN Stalk |
| 220 IgG 1A05 | BN Stalk |
| 241 IgA 2E06 | BN Stalk |

**Table S5: Group 1 HA strains used in Fig. S2C.**

| HA Subtype | Strain Name |
|---|---|
| H1 | A/California/04/2009 |
| H2 | A/Singapore/1/1957 |
| H5 | A/mallard/Italy/3401/2005 |
| H6 | A/chicken/Taiwan/0705/1999 |
| H8 | A/turkey/Ontario/6118/1968 |
| H9 | A/swine/HongKong/9/1998 |
| H11 | A/duck/England/1/1956 |
| H12 | A/duck/Alberta/60/1976 |
| H13 | A/gull/Maryland/704/1977 |
| H16 | A/black-headed-gull/Turkmenistan/13/1976 |
| H17 | A/little-yellow-shouldered-bat/Guatemala/060/2010 |
| H18 | A/flat-facedbat/Peru/033/2010 |

**References**


Cotter, C.R., Jin, H., and Chen, Z. (2014). A single amino acid in the stalk region of the H1N1pdm influenza virus HA protein affects viral fusion, stability and infectivity. PLoS Pathog *10*, e1003831.

Doud, M.B., Lee, J.M., and Bloom, J.D. (2018). How single mutations affect viral escape from broad and narrow antibodies to H1 influenza hemagglutinin. Nat Commun *9*, 1386.

Dreyfus, C., Laursen, N.S., Kwaks, T., Zuijdgeest, D., Khayat, R., Ekiert, D.C., Lee, J.H., Metlagel, Z., Bujny, M.V., Jongeneelen, M.*, et al.* (2012). Highly conserved protective epitopes on influenza B viruses. Science *337*, 1343-1348.

Fu, Y., Zhang, Z., Sheehan, J., Avnir, Y., Ridenour, C., Sachnik, T., Sun, J., Hossain, M.J., Chen, L.M., Zhu, Q.*, et al.* (2016). A broadly neutralizing anti-influenza antibody reveals ongoing capacity of haemagglutinin-specific memory B cells to evolve. Nat Commun *7*, 12780.

Lang, S., Xie, J., Zhu, X., Wu, N.C., Lerner, R.A., and Wilson, I.A. (2017). Antibody 27F3 Broadly Targets Influenza A Group 1 and 2 Hemagglutinins through a Further Variation in VH1-69 Antibody Orientation on the HA Stem. Cell Rep *20*, 2935-2943.

Lingwood, D., McTamney, P.M., Yassine, H.M., Whittle, J.R., Guo, X., Boyington, J.C., Wei, C.J., and Nabel, G.J. (2012). Structural and genetic basis for development of broadly neutralizing influenza antibodies. Nature *489*, 566-570.

Park, J.K., Xiao, Y., Ramuta, M.D., Rosas, L.A., Fong, S., Matthews, A.M., Freeman, A.D., Gouzoulis, M.A., Batchenkova, N.A., Yang, X.*, et al.* (2020). Pre-existing immunity to influenza virus hemagglutinin stalk might drive selection for antibody-escape mutant viruses in a human challenge model. Nat Med *26*, 1240-1246.

Sui, J., Hwang, W.C., Perez, S., Wei, G., Aird, D., Chen, L.M., Santelli, E., Stec, B., Cadwell, G., Ali, M.*, et al.* (2009). Structural and functional bases for broad-spectrum neutralization of avian and human influenza A viruses. Nat Struct Mol Biol *16*, 265-273.

Tan, G.S., Krammer, F., Eggink, D., Kongchanagul, A., Moran, T.M., and Palese, P. (2012). A pan-H1 anti-hemagglutinin monoclonal antibody with potent broad-spectrum efficacy in vivo. J Virol *86*, 6179-6188.

Vanderlinden, E., Goktas, F., Cesur, Z., Froeyen, M., Reed, M.L., Russell, C.J., Cesur, N., and Naesens, L. (2010). Novel inhibitors of influenza virus fusion: structure-activity relationship and interaction with the viral hemagglutinin. J Virol *84*, 4277-4288.

Wu, N.C., Thompson, A.J., Lee, J.M., Su, W., Arlian, B.M., Xie, J., Lerner, R.A., Yen, H.L., Bloom, J.D., and Wilson, I.A. (2020). Different genetic barriers for resistance to HA stem antibodies in influenza H3 and H1 viruses. Science *368*, 1335-1340.

Xu, R., Zhu, X., McBride, R., Nycholat, C.M., Yu, W., Paulson, J.C., and Wilson, I.A. (2012). Functional balance of the hemagglutinin and neuraminidase activities accompanies the emergence of the 2009 H1N1 influenza pandemic. J Virol *86*, 9221-9232.

Yassine, H.M., McTamney, P.M., Boyington, J.C., Ruckwardt, T.J., Crank, M.C., Smatti, M.K., Ledgerwood, J.E., and Graham, B.S. (2018). Use of Hemagglutinin Stem Probes Demonstrate Prevalence of Broadly Reactive Group 1 Influenza Antibodies in Human Sera. Sci Rep *8*, 8628.
